# The Single Cell Landscape of the Human Vein After Arteriovenous Fistula Creation and Implications for Maturation Failure

**DOI:** 10.1101/2025.01.23.634529

**Authors:** Laisel Martinez, Filipe F. Stoyell-Conti, Marwan Tabbara, Miguel G. Rojas, Simone Pereira-Simon, Nieves Santos Falcon, Reyniel Hernandez Lopez, Daniella Galtes, Christina Kosanovic, Anthony J. Griswold, Xiaofeng Yang, Yan-Ting Shiu, Timmy Lee, Juan C. Duque, Marco A. Ladino, Loay H. Salman, Xiaochun Long, Roberto I. Vazquez-Padron

## Abstract

The biological mechanisms underlying arteriovenous fistula (AVF) maturation in hemodialysis patients remain poorly understood despite decades of research. To address this gap, we investigated the cellular changes in the venous wall after fistula creation in histological biopsies of longitudinal veins and AVF samples (N=23 patients). Using single-cell RNA sequencing of 70,281 cells from pre-access veins, mature, and failed AVFs (N=20 patients), we created a complementary transcriptomic atlas of the human vein before and after anastomosis. Postoperatively, the fistula exhibited increased intimal hyperplasia and cell number but reduced cell density, indicating that extracellular matrix (ECM) deposition was more prominent than cell accumulation. Analysis of 14,475 cells from fistulas obtained within one week of creation revealed that inflammation drives early adaptation across all vascular cell types. This includes the pro-inflammatory activation of endothelial cells (ECs) and production of a hyaluronic acid-rich neointima by fibroblasts. By 13 ± 6 weeks, transcriptomic profiles continue to reflect active healing of the vasculature by ECM-producing myofibroblasts and fibroblasts that were found localized throughout the vascular wall, including the intima, using immunofluorescence and in-situ hybridization. Postoperative ECs maintained significant hemostatic adaptations and upregulation of inflammatory molecules (*ACKR3, ICAM1, IL1R1, COL8A1*) supporting their role as gatekeepers of immune cell infiltration. Comparative analyses of failed versus mature AVFs revealed persistent inflammatory signaling among macrophages, ECs, myofibroblasts, and fibroblasts in association with AVF failure. These findings uncover previously unrecognized cellular and molecular patterns in human veins following AVF creation, providing novel insights and potential therapeutic targets to improve AVF outcomes.

**TRANSLATIONAL STATEMENT:** Arteriovenous fistulas (AVF) are a special type of blood vessels that provides access to a patient’s bloodstream during hemodialysis treatments. The AVF is surgically created by connecting an artery and a vein, after which the vein heals and enlarges in a process called “maturation”. We do not fully understand how maturation occurs. This prevents us from designing therapies that ensure proper enlargement of the AVF, which fails in up to 40% of patients. This study investigates how a vein transforms into an AVF in 43 patients with end-stage kidney disease undergoing surgery for AVF creation. We analyze the modifications of structural components of the vein and of diverse populations of cells that direct healing and maturation. Our findings suggest that, contrary to current beliefs, the best therapies to improve AVF maturation should target the accumulation of non-cellular components in the vein and the inflammatory factors that trigger such accumulation.

## INTRODUCTION

Arteriovenous fistulas (AVF) continue to suffer from a high incidence of maturation failure, resulting in prolonged catheter dependence and increased morbidity of patients.^1,2^ To date, clinical trials aimed at preventing thrombosis or enhancing postoperative reendothelialization have failed to improve AVF maturation outcomes.^3–5^ The benefits of treatments targeting intimal hyperplasia (IH) have also been inconsistent.^6–8^ These setbacks underscore the complexity of the vein’s adaptive response after anastomosis and our limited understanding of postoperative venous remodeling.

We now understand that both the early response of the vein and the changes occurring in the following weeks after anastomosis contribute to maturation or failure.^9,10^ Blood flow and diameter at day 1 postop are predictors of AVF success.^10^ Nonetheless, stenoses can develop at any point during early remodeling of the AVF and are not associated with pre-access vein morphometry.^9^ Histological studies of vein and AVF biopsies from patients undergoing AVF creation in two stages have demonstrated that stenotic remodeling of AVFs is associated with postoperative fibrosis, and only in the presence of high fibrosis is IH significantly associated with maturation failure.^11^ The top activated transcriptional programs after AVF creation are inflammatory signaling and extracellular matrix (ECM) remodeling.^12^ Interestingly, compared with the profound molecular transformation of the vein after anastomosis and arterialization, the transcriptional signatures associated with failure seem subtle.^12,13^ This highlights the need for high-resolution studies of human tissues to find the cells and pathways responsible for AVF stenosis.

In this work, we analyze for the first time the adaptive responses of the human vein at one week and 13 weeks after AVF creation at the single-cell resolution. We define the critical roles of endothelial cells (ECs), myofibroblasts/fibroblasts, and macrophages in driving postoperative inflammation and vascular healing. We provide a novel, cell-specific and human-specific framework for understanding the inflammatory ECM remodeling program after anastomosis. Furthermore, we identify clinically relevant targets associated with AVF maturation failure, opening new avenues for translational research to improve AVF surgery outcomes.

## METHODS

### Study subjects and tissue collection

For single-cell RNA sequencing (scRNA-seq), we enrolled 20 participants ≥21 years of age, with chronic kidney disease stage 5 (CKD5) or end-stage kidney disease (ESKD), and planned AVF creation in two stages at Jackson Memorial Hospital (JMH) or the University of Miami Hospital (UMH) from April 2022 to February 2024 (**Figure 1A, Supplementary Table S1**). From these patients we obtained: 1) six pre-access veins collected at the time of access creation, 2) 12 juxta-anastomotic AVF biopsies (six matured, six failed) collected 91 ± 44 days after creation at the time of transposition, and 3) two rare early AVFs resected within 7 days of fistula creation due to high flow steal syndrome or a planned arteriovenous graft (AVG) extension. For immunofluorescence (IF) analyses, we included longitudinal vein and second-stage AVF samples from 23 additional individuals (11 matured, 12 failed) randomly selected from the University of Miami Vascular Access biorepository (**Supplementary Table S2**). Anatomic maturation failure was defined as an AVF whose transected length did not allow a standard transposition because of stenosis and required a short transposition, AVG extension, or ligation.^12,14^ None of the AVFs underwent endovascular or surgical interventions to assist maturation. The study was performed according to the ethical principles of the Declaration of Helsinki and regulatory requirements at JMH and UMH. The ethics committee and Institutional Review Board at the University of Miami approved the study.

**Figure 1.**
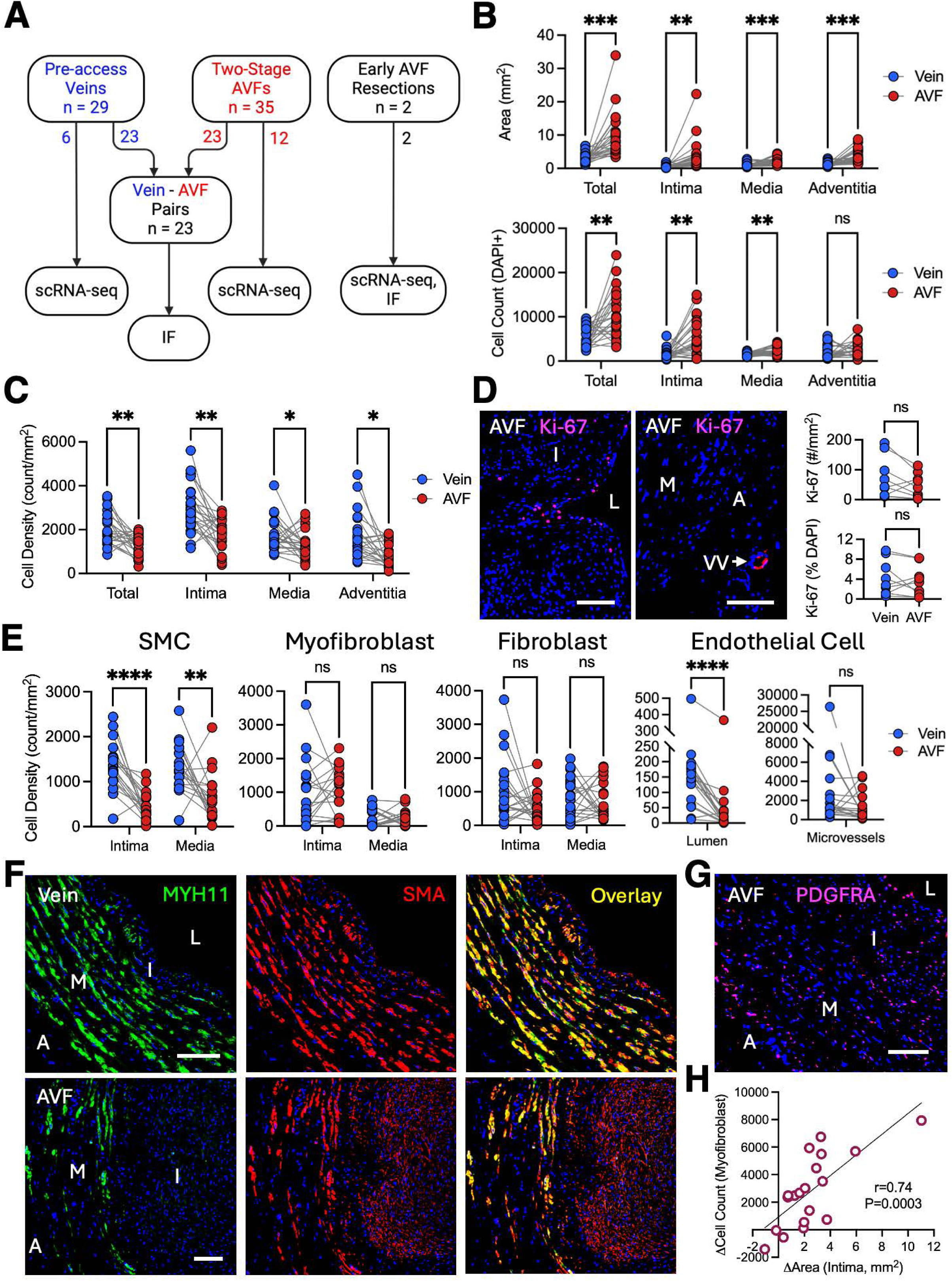
Morphometric and cellular composition changes after arteriovenous fistula (AVF) creation. **A)** Experimental design and tissue collection. **B-C)** Paired comparisons of wall morphometry and cell density in pre-access veins and AVFs. **D)** Immunofluorescence (IF) staining of Ki-67^+^ cells in second-stage AVFs and quantification in tissue pairs. **E)** Cell density of smooth muscle cells (SMC; SMA^+^ MYH11^+^), myofibroblasts (SMA^+^ MYH11^-^), fibroblasts (PDGFRA^+^), and endothelial cells (CD31^+^) in veins and AVFs as quantified by IF. **F)** Representative changes in SMC (yellow in overlay) and myofibroblast (red in overlay) distribution in tissue pairs. **G)** Representative distribution of fibroblasts in second-stage AVFs. **H)** Correlation between the pre-post change in the number of intimal myofibroblasts and the change in intimal area. Image abbreviations: L, lumen; I, intima; M, media; A, adventitia; VV, vasa vasorum. Scale bars = 100 um. *p<0.05, **p<0.01, ***p<0.001, ****p<0.0001

### Single-cell RNA sequencing

Sequencing was performed at the University of Miami John P. Hussman Institute for Human Genomics and analyzed using published bioinformatic pipelines^15–19^ (**Supplementary Methods**). Sequencing data were deposited in the Gene Expression Omnibus, accession number GSEXXXXXX (pending).

### Immunofluorescence and in situ hybridization

Specific cell populations and gene expression markers were validated in tissue sections by IF or in situ hybridization (**Supplementary Methods**).

### Primary cell culture

Human umbilical vein endothelial cells (HUVEC) from Lonza (Walkersville, MD) or primary AVF fibroblasts were stimulated with cytokines or laminal flow as described in the **Supplementary Methods**.

### Statistical analyses

Continuous variables were compared using GraphPad Prism 10.1.1 (San Diego, CA) as indicated in the **Supplementary Methods.**

## RESULTS

### Extracellular matrix deposition dominates over cell proliferation during AVF remodeling

We first evaluated the morphometric and cellular changes in the human vein after anastomosis using pre-access veins and juxta-anastomotic AVF samples from 23 patients undergoing AVF creation in two stages (**Figure 1A**). AVFs experienced significant expansion of the intimal area (5.3 folds [IQR 3.2-9.4]) and moderate growth of the media and adventitia (2.0 [1.2-3.7] and 2.1 [1.3-6.1] folds, respectively) compared with pre-access veins (**Figure 1B**). Surprisingly, despite the higher cell counts in AVFs than veins, the former had fewer cells per mm^2^ (**Figure 1B-C**), suggesting that wall remodeling was dominated by ECM deposition rather than cell accumulation. The reduction in cell density was already apparent in early AVFs (**Supplementary Figure S1A**). Supporting this observation, we found a low proliferative index in AVFs at the time of transposition. The few Ki-67+ cells were predominantly located in the subendothelial region and adventitial microvessels (**Figure 1D**). Cell proliferation was also low one week after surgery (7.6-7.9%; **Supplementary Figure S1B**).

We then explored the changes in cellular composition using IF microscopy and cell type-specific antibodies. AVFs had fewer medial and intimal smooth muscle cells per mm^2^ (SMCs; SMA^+^ MYH11^+^) than pre-access veins, and there was a critical loss of SMCs in early fistulas (**Figure 1E-F** and **Supplementary Figure S1C**). There was also a postoperative reduction of CD31^+^ ECs in the main lumen, suggesting incomplete reendothelialization (**Figure 1E**). In contrast, myofibroblasts (SMA^+^ MYH11^-^) and fibroblasts (PDGFRA^+^) maintained similar densities after AVF creation and were widely distributed throughout the wall (**Figure 1E-G** and **Supplementary Figure S1C**). An expansion of myofibroblasts proportional to the postoperative change in area was confirmed in the intima (r=0.74, **Figure 1H**).

### Single cell sequencing underscores the role of inflammation in the adaptation of the AVF

We constructed a single cell atlas of the human vein before and after anastomosis by RNA sequencing 70,281 unsorted cells from pre-access veins (n=6), early AVF resections (n=2), and second-stage AVFs (n=12). **Figure 2A** presents an unsupervised uniform manifold approximation and projection (UMAP) map with the main cell clusters at resolution 0.5. Vascular populations and immune cells were annotated using canonical gene expression markers previously validated in peripheral veins (**Figure 2B, Data File 1**).^20,21^ As previously reported,^20,21^ there were abundant ECs (18.5 ± 11.5%) due to the high degree of intramural vascularization, fibroblastic cell types (fibroblasts, 18.1 ± 8.9%; myofibroblasts, 9.1 ± 4.2%), mural cells (SMCs, 5.9 ± 6.8%; pericytes, 4.6 ± 4.2%), mono/macs (22.0 ± 12.5%), and NK/T cells (14.6 ± 7.8%). Other immune populations such as mast cells (3.3 ± 2.3%), neutrophils (3.4 ± 5.2%), and B cells (0.6 ± 0.8%) were also found at smaller proportions (**Figure 2A)**. **Supplementary Tables S3** and **S4** present the captured cell proportions in the three types of venous samples.

**Figure 2.**
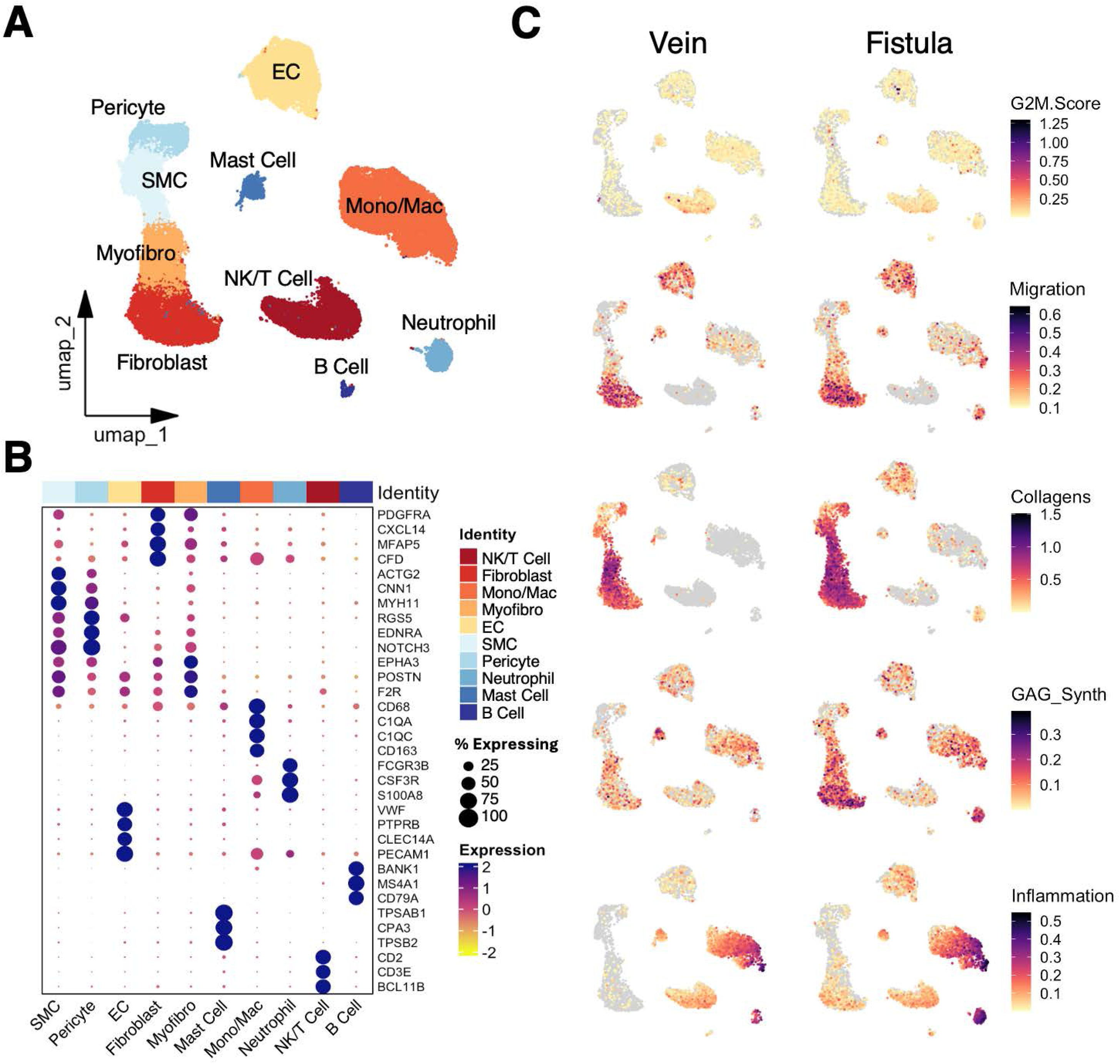
Single cell populations in veins and arteriovenous fistulas (AVFs). **A)** Uniform manifold approximation and projection (UMAP) plot of 70,281 cells from pre-access veins and AVFs. **B)** Dot plot representation of expression markers defining cell populations. Dark blue dots indicate the highest expression levels, while the size of the dot represents the percentage of cells within each cluster expressing the gene as shown in the legend. **C)** Functional profiling of single cell clusters in veins and second-stage AVFs according to gene signature scores. 8,000 cells are projected per tissue type.

Next, we assessed the main remodeling functions of each cell cluster based on transcriptional scores. The inflammatory score (200 genes)^22^ highlighted the role of mono/macs in postoperative inflammation, particularly in the first week after surgery (**Figure 2C** and **Supplementary Figure S2**). Scores of cell cycle genes (S phase: 43 genes; G2/M phase: 54)^23^ indicated a low level of DNA synthesis and cell division in agreement with the Ki-67 stainings (**Figure 1D** and **2C, Supplementary Figure S3**). In contrast, activation of focal adhesion and migration programs (25 genes)^22^ was already evident in myofibroblasts and fibroblasts of early fistulas, as well as in a subset of mono/macs where it likely represents de novo infiltration (**Supplementary Figure S2**). At the time of AVF transposition, fibroblasts and ECs remain the most migratory cell types in AVFs (**Figure 2C**).

In terms of ECM remodeling, almost all vascular populations are involved in the postoperative production of collagens and/or glycosaminoglycans (GAGs) to different extents based on scores that include collagen chains (60 genes) and GAG biosynthetic enzymes (27 genes)^12^ (**Figure 2C** and **Supplementary Figure S2**). Considering that GAGs act as sponge of secretable factors in the ECM, these transcriptional changes agree with the higher inflammatory scores of fibroblasts and myofibroblasts in AVFs compared with veins (**Figure 2C** and **Supplementary Figure S2**). These global changes support a dynamic relationship between inflammation and ECM remodeling that extends beyond the early transformation of AVFs.

### Macrophages are among the first responders to postoperative stress in the AVF wall

To further explore the role of inflammatory cells in the control of venous remodeling, we evaluated the changes in macrophage (CD68^+^) abundance after anastomosis using IF microscopy (**Figure 3A**). We noticed significant infiltration in one-week fistulas and similar density of macrophages in pre-access veins and AVFs (**Supplementary Figure S4A-B**). Re-clustering of the mono/mac population uncovered the effects of AVF creation on the monocytic phenotypes residing the venous wall (**Figure 3B** and **Supplementary Figure S4C**). AVFs were characterized by an increase in ECM-producing macrophages (Mac_4; expressing *COL1A1, COL1A2, DCN, LUM, CCDC80*; 7.8 ± 11.5% vs. 1.2 ± 1.5%, p=0.0064) but a reduction in the tissue-resident phenotype^24^ (Mac_1; expressing *FOLR2, MAMDC2, LILRB5*, *LYVE1*; 25.6 ± 12.4% vs. 52.7 ± 7.9%, p<0.001) (**Figure 3C, Supplementary Tables S5** and **S6, Data File 2**). Inflammatory monocytes (defined by *S100A8, S100A9, AQP9*, and *EREG*) increased 7.5x in second-stage AVFs compared with pre-access veins (11.2 ± 7.2% vs. 1.5 ± 1.4%, p<0.0001), and approximately 20x (23.5 – 36.5%) in one-week AVFs, suggesting de novo infiltration (**Figure 3C, Supplementary Figure S4C, Supplementary Tables S5** and **S6**). On the other hand, lipid-processing macrophages (Mac_2; *TREM2, LIPA, APOE, APOC1*), TGF-β activated (Mac_3; *CD69, CLDN1, CCL3L1, CCL4L2*),^25^ and dendritic cells (DC; *FCER1A, CD1E, CD1C*, *IL1R2*)^26^ had similar proportions in preoperative and postoperative samples.

**Figure 3.**
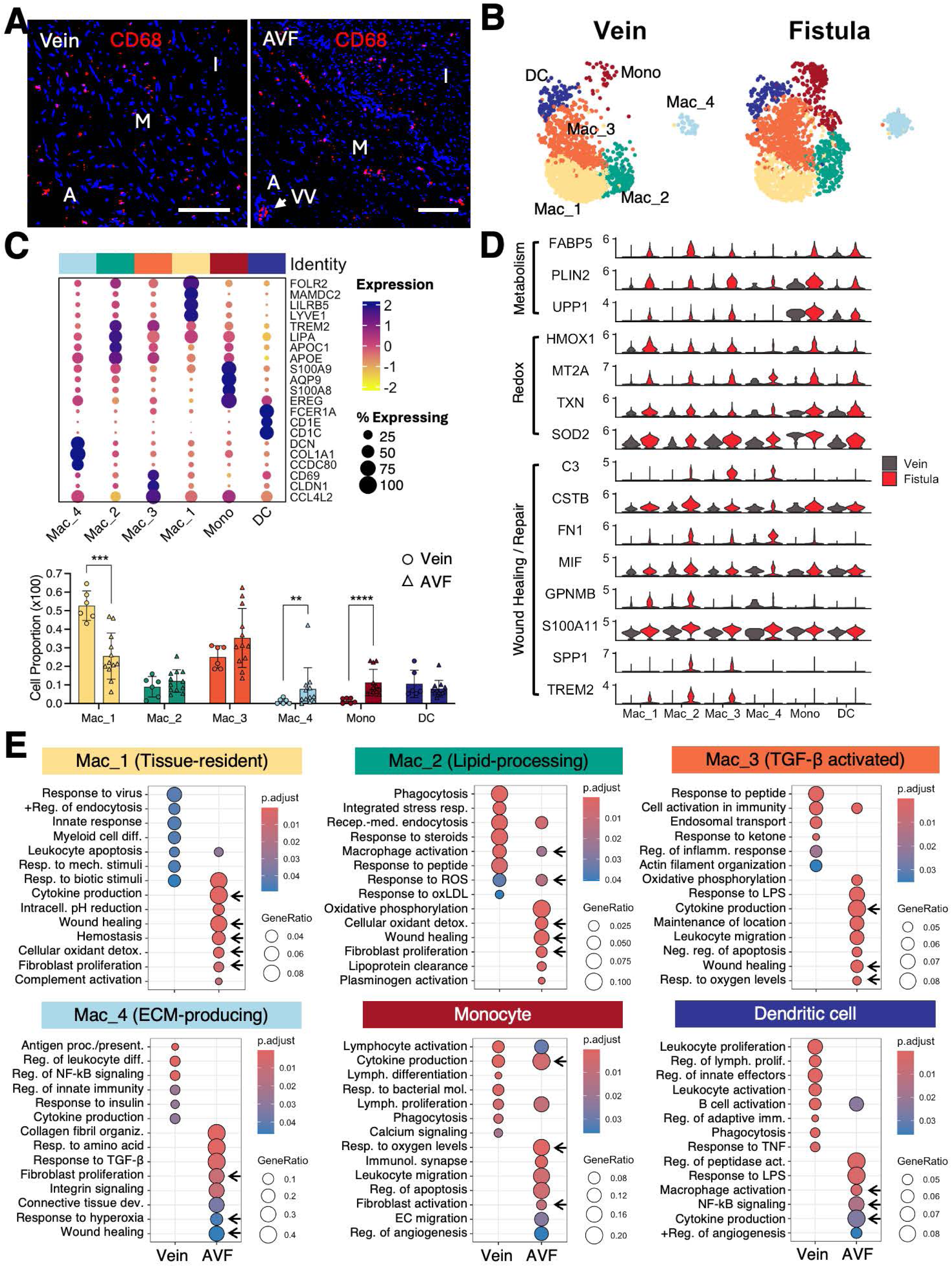
Changes in macrophage phenotypes after arteriovenous fistula (AVF) creation. **A)** Representative distribution of macrophages in vein and AVF pairs. I, intima; M, media; A, adventitia; VV, vasa vasorum. Scale bars = 100 um. **B)** Focused UMAPs of monocytic phenotypes in pre-access veins and second-stage AVFs. 2,000 cells are projected per tissue type. **C)** Dot plot representation of expression markers defining mono/mac subclusters, and proportions of phenotypes relative to the overall cluster in veins and second-stage AVFs. **p<0.01, ***p<0.001, ****p<0.0001 **D)** Violin plots of selected expression changes after AVF creation. **E)** Gene ontology (GO) over-representation analysis of biological processes activated in veins and second-stage AVFs. Arrows indicate processes in common among postoperative phenotypes related to hemodynamic adaptations and vascular repair.

Analysis of gene expression changes in the overall mono/mac cluster after fistula creation detected 680 differentially expressed genes (DEGs; avg_log_2_FC>|1|, p_val_adj<0.01) between second-stage AVFs and pre-access veins (**Data File 3**). Interestingly, the list of DEGs reflected adaptive changes associated with oxidative stress responses, fibroblast activation, and participation in wall repair (**Figure 3D-E** and **Supplementary Figure S5B**). Most mono/mac phenotypes in AVFs showed a pro-inflammatory metabolic signature characterized by high expression of *FABP5, PLIN2,* and *UPP1*.^27–29^ They also showed upregulation of redox enzymes and chelators (*HMOX1, MT2A, TXN, SOD2*) in response to heme and reactive oxygen species (ROS). Upregulated genes involved in wound healing included *SPP1, S100A11, FN1*, and *GPNMB*, which activate fibroblasts and tissue remodeling.^30–32^ Expression of *TREM2* and *C3* in macrophages is also associated with better clearance of necrotic cells and debris.^33,34^ Comparison with early AVFs indicates that these adaptive changes occur as early as the first week after anastomosis (**Supplementary Figure S5B**), highlighting the role of myeloid cells in venous hemostasis and wall repair.

### Reparative EC phenotypes emerge in AVFs in response to flow and inflammation

We then examined the effects of AVF creation on ECs and their potential relationship with infiltrating monocytes. We found 5 distinct EC subtypes in preoperative and postoperative tissues (**Figure 4A-D**). EC_1 cells are mostly of a “venous” phenotype (expressing *PLVAP, FABP4, COL15A1*, *ZNF385D*, *ACKR1*) but also contain small subsets of arteriolar and capillary ECs that did not separate from the main subcluster (**Supplementary Figure S6A**). EC_2 cells have a valvular-like phenotype (*EFEMP1, BMP4, OMD*, *PLXNA4, CRTAC1, GDF7*), while EC_3s have lymphatic characteristics (*TFF3, SCG3, PROX1*, *FOXC2*, *STXBP6*) (**Figure 4A-B, Data File 4**). The EC_4 phenotype is defined by shear stress genes and hemostatic modulators (*PI16, ECM1, SERPINE1, SERPINE2, PTGIS, PTGDS*). EC_5 cells, in turn, represent the end stage of endothelial-mesenchymal transition (endoMT; expressing *COL1A1, COL1A2, DCN, COL6A3*). Interestingly, unlike EC_1 cells, which express the typical venous marker ACKR1^20^ (**Supplementary Figure S6A**), the rest of the EC phenotypes show high expression of the chemokine receptor *ACKR3* (**Figure 4C**). Among these, the hemostatic EC_4 cells are 8x more abundant in second-stage AVFs compared with pre-access veins (19.8 ± 20.6% vs. 2.4 ± 3.6%, p=0.0057), while EC_2 cells demonstrate a significant reduction (4.5 ± 3.5% vs. 20.0 ± 9.6%, p=0.0033) (**Figure 4D, Supplementary Table S7**). These changes are early adaptations already present in the first week after anastomosis (**Supplementary Table S8**). ACKR3 stainings in pre-access veins and AVFs indicate that, while some veins have ACKR3^+^ cells in the main lumen, the increasing EC_4 population in AVFs is predominantly located in the adventitia (**Figure 4E** and **Supplementary Figure S6B**).

**Figure 4.**
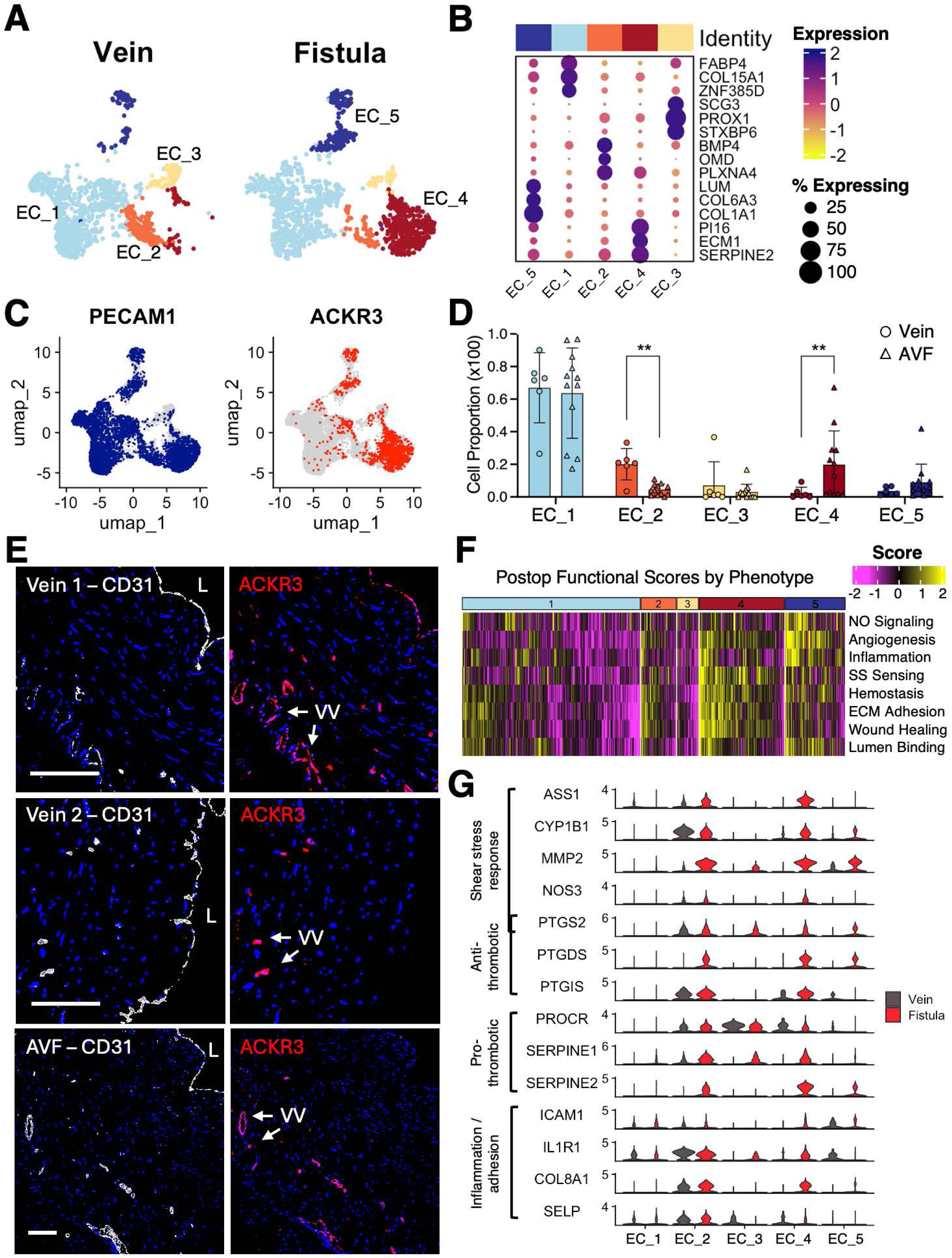
Functional adaptations of endothelial cells (EC) after arteriovenous fistula (AVF) creation. **A)** Focused UMAPs of EC phenotypes in pre-access veins and second-stage AVFs. 1,500 cells are projected per tissue type. **B)** Dot plot representation of expression markers defining EC subclusters. **C)** Expression of CD31 in the overall cluster and of the atypical chemokine receptor 3 (ACKR3) in selected phenotypes. **D)** Proportions of EC phenotypes relative to the overall cluster in veins and second-stage AVFs. **p<0.01 **E)** Representative distribution of ACKR3^+^ ECs in veins and AVFs. ACKR3^+^ ECs may be present (Vein 1) or absent (Vein 2) in the main lumen of veins but are predominantly found in the vasa vasorum of AVFs. L, lumen; VV, vasa vasorum. Scale bars = 100 um. **F)** Heatmap of functional profiles of postoperative ECs based on gene signature scores. Yellow indicates activation of processes. **G)** Violin plots of adaptive gene expression changes after AVF creation.

Analysis of gene expression changes in the overall EC cluster in AVFs vs. veins uncovered 692 DEGs, including postoperative upregulation of genes involved in hemostasis (*PTGDS, SERPINE1, SERPINE2*), permeability (*SPOCK1*), inflammation (*ACKR3, BMP2*), ECM remodeling (*COL4A1, COL1A1, MMP2, PI16*), and interaction with fibroblasts (*CLIC3, MFAP5*) (**Supplementary Figure S7, Data File 5**). Scoring of cells using annotated pathways in the Molecular Signatures Database (MSigDB^22^; **Supplementary Methods**) identified the EC_4 population as the main subtype responsible for adaptive shear stress sensing, hemostasis, and wound healing mechanisms (**Figure 4F**). Specifically, EC_4 cells showed enrichment of shear stress response genes such as *ASS1, CYP1B1, MMP2, NOS3*, and *PTGS2* (also known as COX-2); anti-thrombotic (e.g., *PTGS2, PTGDS, PTGIS*) and pro-thrombotic mediators (e.g., *PROCR, SERPINE1, SERPINE2*); as well as inflammatory receptors and adhesion molecules (e.g., *ICAM1, IL1R1, COL8A1, SELP*; **Figure 4F-G**) as early as one week after anastomosis (**Supplementary Figure S8**). These genes were also upregulated in the EC_2 cells that decrease after anastomosis (**Figure 4G**), suggesting these subpopulations are phenotypically related.

We tested the contribution of inflammatory stimuli and laminar flow to the postoperative programs of ECs (**Figure 5**). Treatment of HUVEC cells revealed that IL-1β induced inflammatory receptors, hemostatic regulators, and basement collagens (*ACKR3, COL8A1, PTGIS, PTGS2*), but not shear stress response transcripts (*ASS1, CYP1B1, MMP2, NOS3*) (**Figure 5A-B**). In contrast, exposure to flow induced shear stress response genes, *PTGIS*, *PTGS2*, and *COL4A1*, but not *ACKR3* and *COL8A1* (**Figure 5C**). Treatment with an NF-kB inhibitor or a p65-targeted siRNA reduced the induction of *ACKR3* and *COL8A1* by IL-1β (**Figure 5D-E**). These results indicate that postoperative transcriptional programs in ECs are the results of hemodynamic and inflammatory effects.

**Figure 5.**
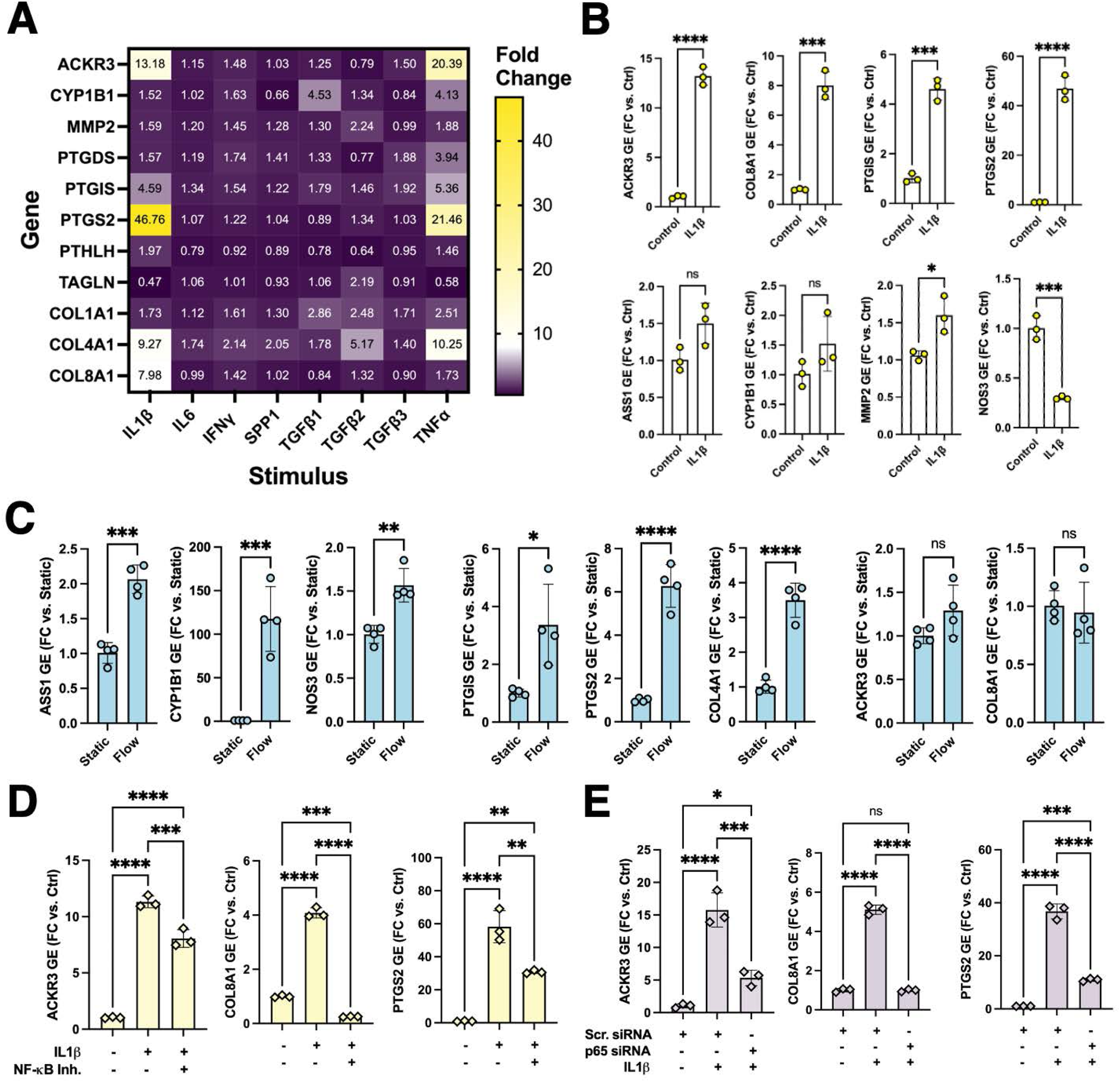
Transcriptional regulation of postoperative EC characteristics. **A)** Heatmap of fold change induction in the RNA expression of inflammatory, hemostatic, shear sensing, and extracellular matrix genes in HUVEC cells treated with inflammatory stimuli. **B)** IL-1β induces inflammatory and hemostatic genes, downregulates eNOS, but has minor to no effects on other shear stress genes. **C)** Laminal flow induces hemostatic and shear stress genes, but not ACKR3 or COL8A1. **D-E)** Induction of inflammatory and hemostatic genes by IL-1β is reduced by pre-treatment with NF-κB inhibitors and RelA/p65 targeted siRNAs. Fold changes are reported as mean ± SD relative to the control or scrambled siRNA treatment. *p<0.05, **p<0.01, ***p<0.001, ****p<0.0001

### Fibroblasts transform the ECM of the neointima and participate in wall repair

We then took advantage of scRNA-seq analyses to better define the postoperative fate of mural cells and fibroblast/myofibroblasts after anastomosis (**Supplementary Figure S9A**). As expected, SMCs shifted towards a synthetic phenotype, showing postoperative upregulation of *FN1, COL8A1, IGFBP2, POSTN*, and *FAP*, as well as lower expression of contractile genes (*MYH11, DES*) (**Supplementary Figure S9B, Data File 6**). Both SMCs and pericytes from AVFs displayed increased expression of ECM proteins (e.g., *COL1A1, COL3A1, LUM, VCAN, OGN*) and hypoxia-inducible genes (e.g., *HIGD1B, NDUFA4L2*) compared with the corresponding cells in veins (**Supplementary Figure S9B-C, Data Files 6-7**). Postoperative pericytes also showed upregulation of PLXDC1, an important receptor for angiogenesis.^35^

Venous fibroblasts (F; expressing *CFD, C3*, *APOD*) grouped in four phenotypes while myofibroblasts (MF; *MYH11^-^*, *DES^-^, ACTA2^+^, POSTN^+^*, *F2R^+^*)^21^ formed two subclusters (**Figure 6A-C**, **Supplementary Figure S10, Data File 8**). AVF creation significantly increased the proportions of healing fibroblasts (F_3 and F_4), decreased regulatory phenotypes (MF_2 and F_1), and further stimulated activated populations (MF_1 and F_2) (**Figure 6B, Supplementary Tables S9** and **S10**). Inflammatory F_4 fibroblasts (expressing *IL1B, CXCL8*) were the first to appear in early fistulas likely in response to inflammation and/or hemodynamic or vascular injury. They were also the first to express hyaluronan synthase 1 (*HAS1*) to form a provisional ECM^36^ and may be a precursor population to *HAS1* and *HAS2-*positive F_3 cells in second-stage AVFs (**Figure 6C, Supplementary Figure S10A**). The F_3 phenotype increased almost 8x in AVFs compared to pre-access veins (2.7 ± 1.9% vs. 21.3 ± 16.9%, p<0.001), suggesting a neointimal localization. Their widespread distribution and accumulation in the neointima were confirmed by HAS1 IF in two-stage fistulas (**Figure 6D**). Many of them also expressed POSTN indicating a state of activation. MF_2 cells, which regulate ECM remodeling and fibroblast activation^37–39^ (*COMP, PRELP, LTBP2*), as well as F_1 fibroblasts that control detoxification and metabolism (*ADH1B, ABCA10*) decreased by approximately one half in AVFs (MF_2, 22.2 ± 5.1% vs. 11.5 ± 8.3%, p=0.019; trend in F_1, 26.9 ± 9.5% vs. 15.0 ± 10.7%, p=0.054). Two activated phenotypes, MF_1 (expressing *ACTA2, AGT, POSTN*) and F_2 (DPP4, *SEMA3C, WNT*),^40–42^ did not change in proportions after in two-stage AVFs compared with veins, but did demonstrate postoperative upregulation of mechanosensitive genes (e.g., *THBS1, PIEZO2, THY1*)^43^ and pro-fibrotic signals (*COL1A1, COL1A2, COL3A1, POSTN, SPARC, TGFBI*) (**Supplementary Figure S11**). The POSTN^+^ HAS1^-^ MF_1 population was also found in the postoperative intima by IF/ISH (**Figure 6D**).

**Figure 6.**
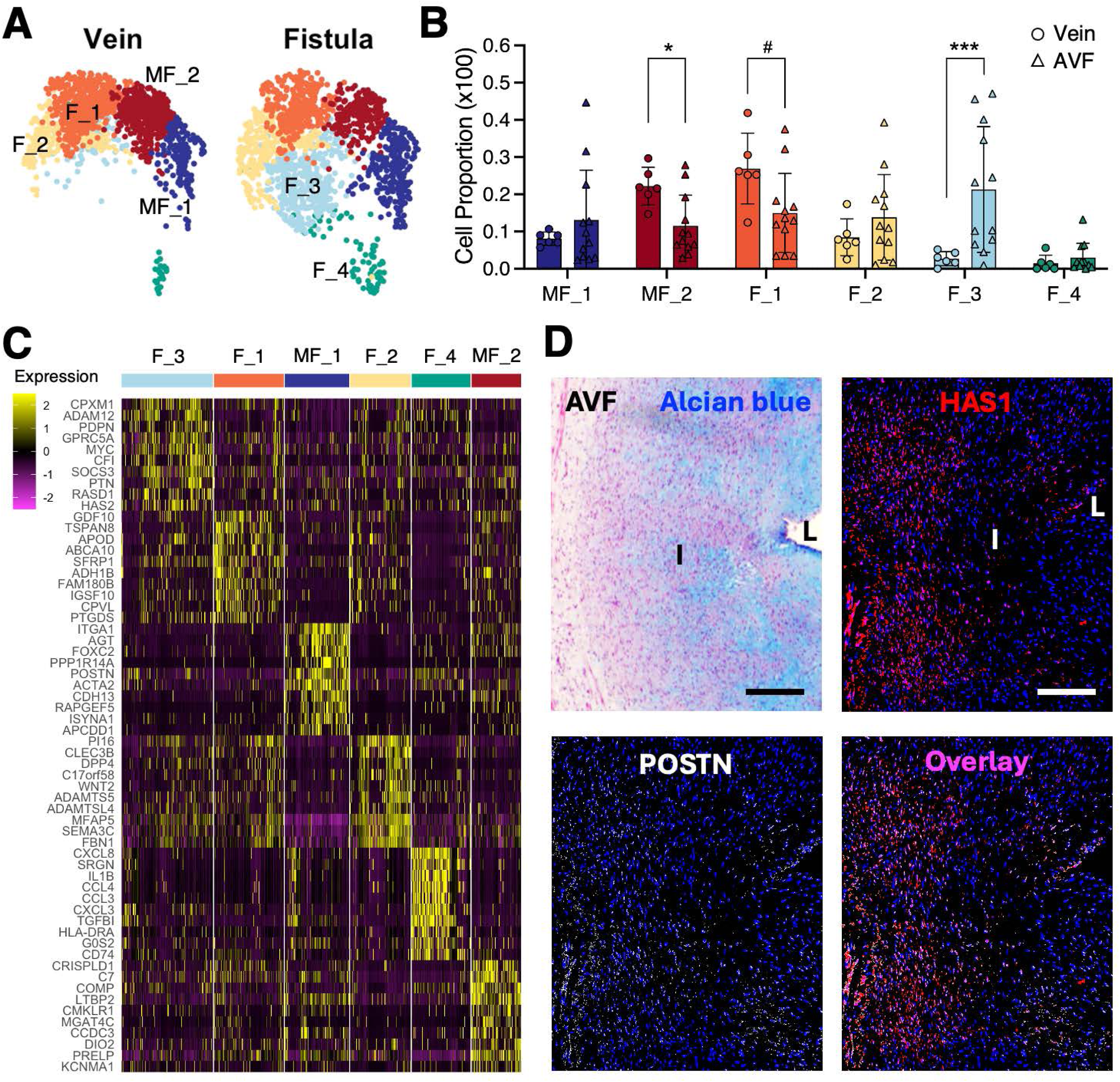
Myofibroblasts and fibroblasts after arteriovenous fistula (AVF) creation. **A)** Focused UMAPs of myofibroblast (MF) and fibroblast (F) phenotypes in pre-access veins and second-stage AVFs. 1,500 cells are projected per tissue type. **B)** Proportions of MF and F phenotypes relative to the overall cluster in veins and second-stage AVFs. *p<0.05, **p<0.01, #p=0.054 **C)** Heatmap representation of expression markers defining subpopulations of myofibroblasts and fibroblasts. Upregulation is shown in yellow. **D)** Representative localization of F_3 fibroblasts (HAS1^+^, red or pink in overlay) and MF_1 myofibroblasts (POSTN^+^ HAS1^-^, white in overlay) in second-stage AVFs, and corresponding Alcian blue staining. L, lumen; I, intima; M, media; A, adventitia. Scale bars = 200 um.

Gene ontology enrichment analyses suggest that all myofibroblast and fibroblast subtypes participate in AVF remodeling, albeit through different mechanisms (**Figure 7A**). Not surprisingly, we found transcriptional overlaps among the processes of wound healing, tissue regeneration, and mechanical responses (**Figure 7B**). In MF_1 cells, the processes involve upregulation of integrin components (*ITGA2, ITGB1, ILK, PTK2, TLN1, VCL*), pro-angiogenic factors (*ANGPT2, TNC*), ECM remodeling proteins (*ENG, FLNA, FN1, MATN2, POSTN, TGFB1, TIMP1*), and hemostatic genes (*F2R, SERPINE1*) (**Data File 8**). The F_4 population from early AVFs participates in wound healing by regulating angiogenic responses (*HIF1A*), cell-ECM adhesion (*CLIC1, ACTB, ACTG1, CNN2*), clotting factor production (*LMAN1, VKORC1*), and activation of fibroblasts (*FIBP*). Lastly, tissue regeneration and mechanosensing by the neointimal F_3 phenotype involves mitogenic and pro-angiogenic factors (*MDK, PTN, ADM*), hemostatic genes (*SERPINE2*), and mechano-regulators such as *CITED2*^44^ and *THBS1* (**Data File 8**).

**Figure 7.**
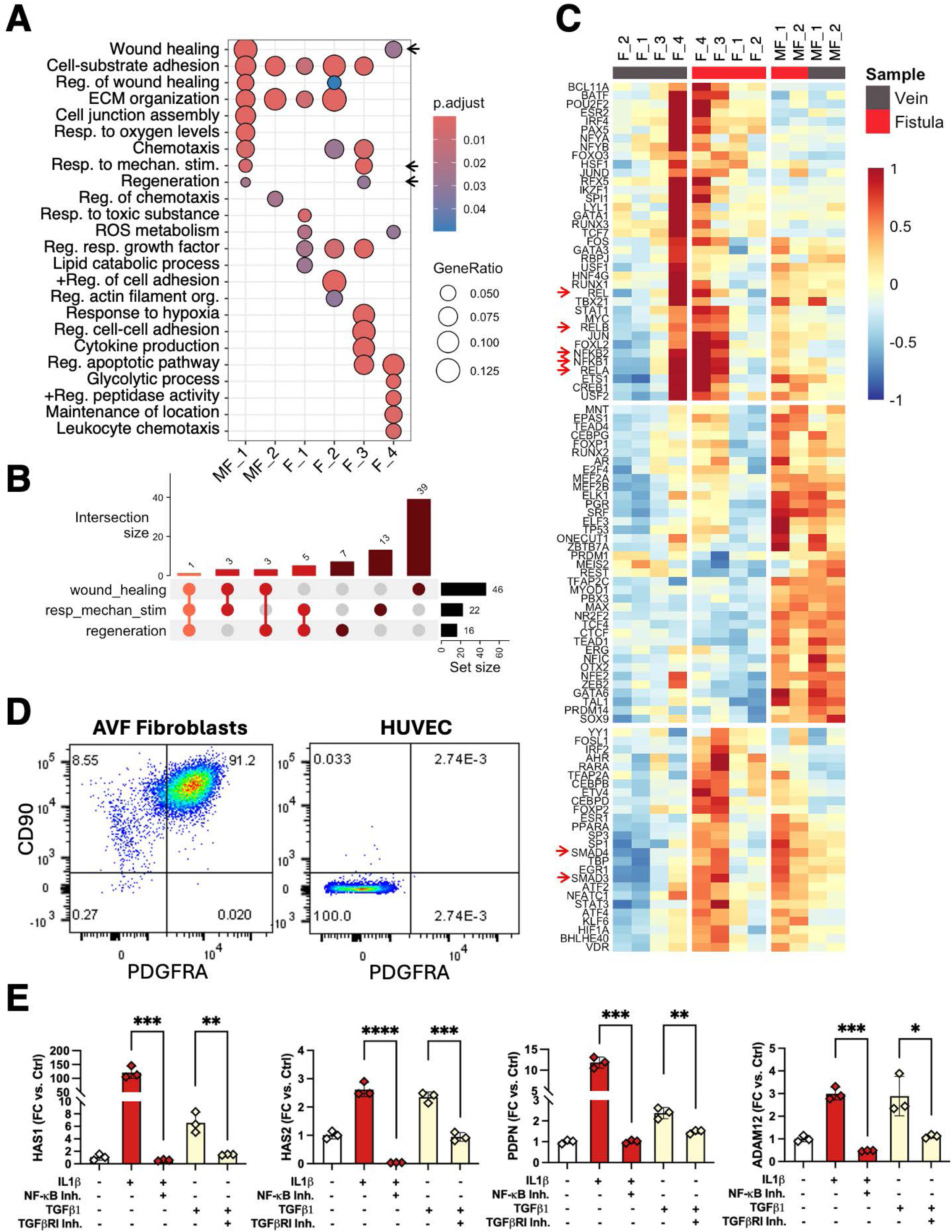
Transcriptional regulation of postoperative fibroblast differentiation. **A)** Gene ontology (GO) over-representation analysis of biological processes activated in myofibroblast and fibroblast subtypes. Arrows indicate mechanosensitive and vascular healing pathways. **B)** Functional overlap among the GO processes of wound healing, tissue regeneration, and responses to mechanosensitive stimuli based on the number of shared genes in myofibroblasts and fibroblasts. **C)** Prediction of top 50 transcription factors in myofibroblasts and fibroblasts from veins and AVFs. Arrows point to members of the NF-κB and SMAD families. **D)** Flow cytometry analysis of PDGFRA and CD90 (THY-1) in primary human AVF fibroblasts and control cells (HUVEC). **E)** Treatment of human AVF fibroblasts with IL-1β and TGF-β1 induce the RNA expression of genes characteristics of healing fibroblasts. The effects are reduced by pre-incubation with NF-κB and TGFβRI inhibitors, respectively. Fold changes are reported as mean ± SD relative to the control treatment. *p<0.05, **p<0.01, ***p<0.001, ****p<0.0001

To understand the differentiation of the postoperative fibroblast phenotypes, we predicted the top 50 transcription factors activated in myofibroblasts and fibroblasts from veins and AVFs based on the expression of target genes in the DoRothEA database.^45^ This analysis demonstrated that activation of NF-kB is a defining feature of the neointimal F_3 fibroblasts and the F_4 population (**Figure 7C**), the most directly involved in vascular repair. Increased activity of SMAD3 and 4, which are part of the TGF-β signaling pathway, is also observed in postoperative myofibroblasts as well as F_3 and F_4 cells. Using primary fibroblasts from human AVFs (**Figure 7B**), we further demonstrated the induction of genes characteristic of the F_3 population (*e.g., HAS1, HAS2, ADAM12, PDPN*) by IL-1β and TGF-β1 to a lower extent. The effect was abrogated by pre-incubation with NF-kB and TGFβRI inhibitors. (**Figure 7C**). These results indicate that inflammation is a critical trigger for the differentiation of AVF fibroblasts.

### Single-cell analyses uncover an inflammatory hub among mono/macs, ECs, and fibroblasts associated with AVF failure

We performed global cell-to-cell communication analyses using CellChat v2^19^ to infer the cell interactions that most contributed to maturation failure (**Figure 8A-B**). A higher number and strength of interactions was predicted in AVFs that failed compared with those that matured (**Figure 8A**), mainly due to enhanced communications among mono/macs, ECs, fibroblasts, and myofibroblasts (**Figure 8B** and **Supplementary Figure S12A**). Supporting this inflammatory hub, we found a 2x higher trend of mono/macs in AVFs that failed compared with mature fistulas (25.6 ± 11.1% vs. 12.6 ± 5.3%, p=0.054) by scRNA-seq (**Figure 8C, Supplementary Table S11**), which was confirmed by CD68^+^ IF in the media of AVFs (**Figure 8D**).

**Figure 8.**
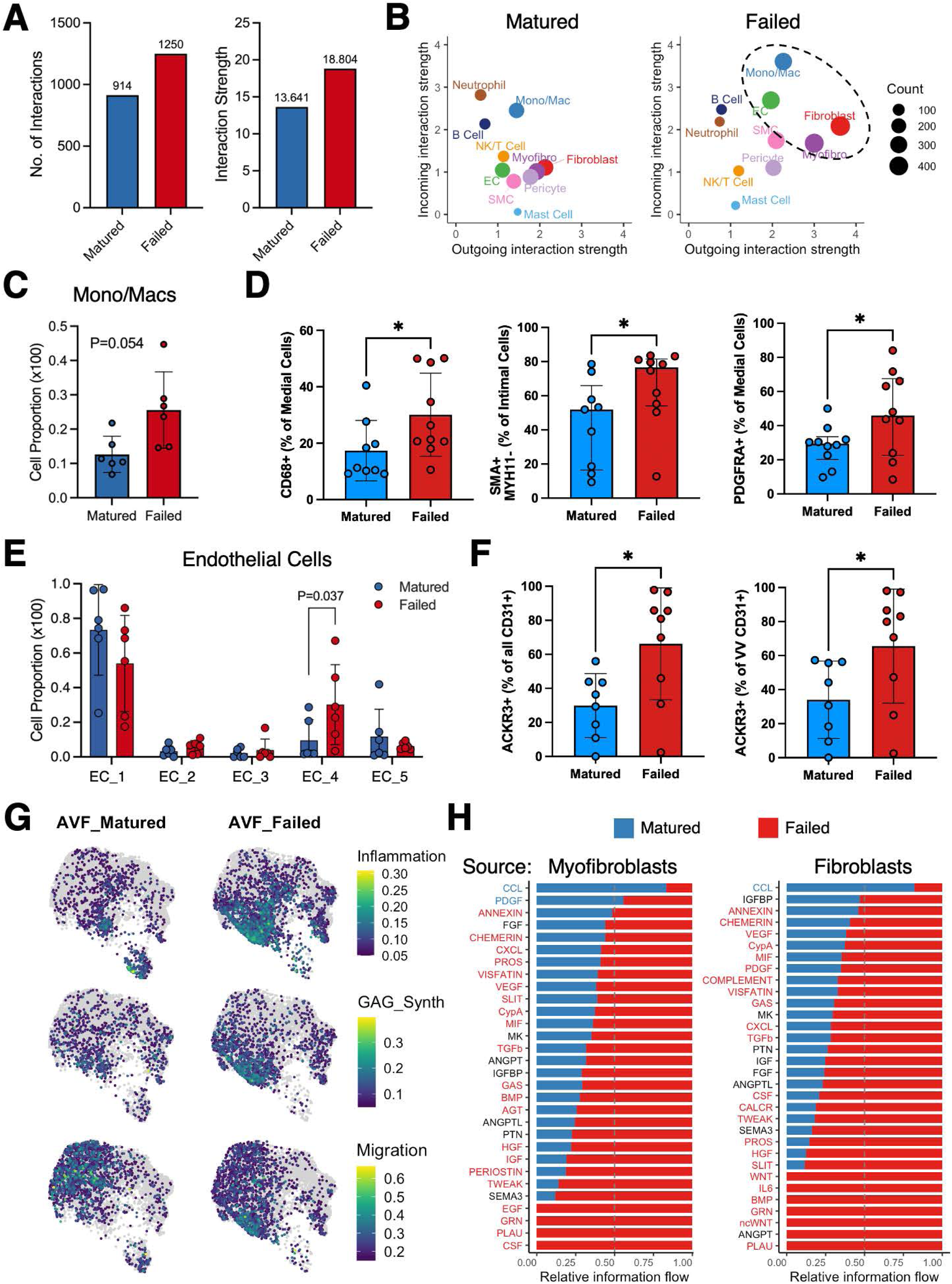
Single cell signatures associated with maturation failure. **A)** Number and strength (probability) of cell-to-cell interactions in AVFs that matured and failed as predicted by CellChat. **B)** Enhanced cross-talk of endothelial cells (EC), myofibroblasts, fibroblasts, and mono/macs in AVFs that failed. The sizes of circles indicate the number of predicted interactions. **C)** Increasing trend in the proportion of mono/macs in AVFs that failed compared to those that matured by scRNA-seq. **D)** Differences in cell compositions between AVFs that matured and failed by immunofluorescence (IF). **E-F)** Increased single cell abundance of inflammatory-hemostatic ECs in association with maturation failure and confirmed by IF in independent AVFs. Bar graphs indicate mean ± SD or median ± IQR as appropriate. *p<0.05 **G)** Functional profiles of myofibroblasts and fibroblasts from AVFs that matured and failed based on gene signature scores. 8,000 cells are projected per tissue type. **H)** Cell-to-cell communication pathways in AVFs that matured and failed using myofibroblasts and fibroblasts as sources of ligands. Pathways with blue or red font are significantly associated with maturation or failure, respectively.

Comparisons of EC subtypes also demonstrated significant differences between failed and mature AVFs. Inflammatory-hemostatic ECs (EC_4) had significantly higher proportion in failed fistulas (30.2 ± 23.1% vs. 9.5 ± 12.1%, p=0.037), as confirmed by ACKR3^+^ IF (**Figure 8E-F, Supplementary Table S13**). As a result, genes enriched in this population had the highest fold change and statistical support in association with AVF failure. These included receptors of the IL-1 family (*IL1R1, IL1RL1*), the chemerin chemokine-like receptor 1 (*CMKLR1*), and *COL8A1*, which was previously identified as a risk factor for failure by bulk RNA-seq^12^ (**Data File 9**).

Other differences in cell composition associated with maturation failure include more medial PDGFRA^+^ fibroblasts and more intimal SMA^+^ MYH11^-^ myofibroblasts independent of intimal area (2.90 [IQR 1.20-6.64] vs. 2.03 [1.43-3.24] mm^2^ in mature vs. failure, respectively; p=0.24) (**Figure 8D**). In addition, neointimal fibroblasts (F_3) from AVFs that failed had higher scores of inflammation, GAG synthesis, and migration (**Figure 8G**). Increased inflammatory and oxidative stress in myofibroblasts and fibroblasts in association with AVF failure is evidenced by the upregulation of phospholipase C gamma 2 (*PLCG2*), *PLA2G2A*, as well as metallothioneins (e.g., *MT2A, MT1E, MT1X*) and antioxidant genes (e.g., *HMOX1, TXN*) in multiple phenotypes (**Supplementary Figures S13** and **S14**). Interestingly, many of the failure-associated DEGs detected in ECs were also upregulated with failure in these populations. These included *COL8A1*, several hemostatic genes (*SERPINE1, THBD*), as well as inflammatory factors (*CXCL1, CXCL2, IL6, LIF*) and receptors (*IL1R1, IL1RL1*) (**Supplementary Figures S13** and **S14, Data Files 10-17**). This inflammatory microenvironment may be supported by the GAG enzymes *B3GNT2, CHST11,* and *ST3GAL1*, which have higher levels in neointimal F_3 fibroblasts of failed AVFs (**Supplementary Figure S14**).

Finally, using cell-to-cell interactome analyses, we interrogated the signaling pathways that used myofibroblasts or fibroblasts as sources of ligands, directed toward mono/macs, ECs myofibroblasts, and fibroblasts as receptors (**Figure 8H**). Only two pathway interactions were enhanced in mature AVFs (CCL, PDGF) while more than 20 were significantly enriched in AVF failure (e.g., AGT, BMP, CSF, CXCL, MIF, PLAU). This analysis supports a unique pro-inflammatory microenvironment in AVFs that failed that prevents inflammation resolution during postoperative wall repair.

## DISCUSSION

This work presents for the first time the transcriptional profiles of >70,000 individual cells from pre-access human veins and AVFs and provides an opportunity to explore the intrinsic mechanisms underlying AVF remodeling without the limitations of animal models. Our findings highlight: 1) the critical role of early inflammation in shaping postoperative cellular phenotypes, 2) the fibrotic nature of venous arterialization, 3) the adaptive capacity of adventitial ECs in AVFs, and 4) the contribution of fibroblasts to sustained inflammation through modification of ECM composition. Additionally, we identify cellular patterns associated with AVF failure that hold significant potential for translational research.

We studied the single-cell transcriptomes of rare early resections obtained 5-7 days after anastomosis. This timeframe is particularly relevant, since early adaptation of the vein has been shown to predict AVF outcomes.^10^ We confirmed the significant infiltration of monocytes, quick activation of hemostatic genes in ECs, and phenotypic modulation or loss of SMCs in the first week postop, as previously reported.^46^ We also discovered a paradoxical low proliferation index (<10%) in the AVF regardless of the collection time and thickening of the wall. Therefore, either cell proliferation went uncovered by the limited time points included in the study or AVF remodeling is mostly driven by cell mobilization and ECM deposition. This agrees with previous studies of proliferation index in AVFs^47,48^ and with the significantly lower density of SMCs in AVFs compared with pre-access veins. To date, research on AVF stenosis has been mostly focused on cell proliferation, which has translated into the local delivery of cytostatic drugs with suboptimal success.^6^ The results from ongoing trials will further clarify the benefit of these treatments for improving AVF maturation in patients aged 65 and older.^49^

This study also emphasizes the essential role of inflammation in the adaptation of the vein after fistula creation. It is now evident that inflammation characterizes early cellular phenotypes of both immune and non-immune origin, but gradually resolves over time. Notably, our findings show that the inability to resolve inflammation in certain cells can contribute to maturation failure. For example, while inflammatory fibroblast phenotypes typically subside in mature AVFs, this resolution is less evident in cases of failure, where a signaling hub among immune and non-immune cells is clearly established. In this hub, ECs, myofibroblasts and fibroblasts are not only the receivers (receptors) but also senders of multiple inflammatory signals (ligands). This complex environment may underlie the failure of the recent prednisolone trial in AVF maturation,^50^ since glucocorticoids can enhance TGF-β signaling and fibroblast activation.^51^ Persistent inflammation may also underlie the excessive fibrosis that is associated with failure, as prolonged exposure to cytokines (e.g., IL-1β) is known to increase SMAD3 phosphorylation and enhance TGF-β signaling.^52^

We further showed a postoperative reduction of CD31^+^ ECs in the main lumen, but resilient EC populations in the vasa vasorum that adapted phenotypically to the new hemodynamic conditions. The loss of luminal CD31^+^ cells may reflect a replacement by EC-derived mesenchymal cells or a true incomplete reendothelialization. Nonetheless, these changes do not seem to affect AVF maturation, likely due to the supra-arterial blood flow that prevents thrombosis and the anticoagulant properties of GAGs in the neointima.^53^ In the case of the adventitial ECs, their postoperative adaptation included the activation of shear stress sensing pathways possibly due to retrograde flow from the lumen,^54^ pro-inflammatory features, and hemostatic mechanisms. These adaptive changes established the inflammatory status that influenced AVF maturation outcomes. An increase in ACKR3^+^ ECs was observed in unsuccessful AVFs, in parallel with higher infiltration of mono/macs. ACKR3 plays a role in regulating the intramural concentration of chemokines such as MIF and CXCL12, which induce leukocyte arrest and transmigration.^55,56^ The expression of ACKR3 is induced by inflammatory factors like IL-1β as demonstrated in our study. This represents yet another inflammatory loop that remains active in AVF failure and could potentially be targeted in the future to improve AVF outcomes.

Finally, our results highlight a heterogeneous population of fibroblasts and myofibroblasts as central players in postoperative venous remodeling. These cells undergo significant reprogramming in response to mechanical stimuli (e.g., over-stretching), changes in ECM composition, and immune ligands. Gene expression profiles indicate that fibroblasts and myofibroblasts contribute to hemostasis by continuously repairing the fistula wall. Notably, fibroblasts appear to be the primary source of the hyaluronic acid-rich matrix that characterizes the neointimal layer of the AVF. As ACKR3^+^ ECs, this matrix is also induced by inflammatory signals, and may have additional GAG features (e.g., upregulation of *B3GNT2, CHST11,* and *ST3GAL1*) that correlate with sustained inflammation in AVF failure. Considering the various potential origins for neointimal myofibroblasts^57^ and the innumerable DEGs between myofibroblasts and fibroblasts from mature and failed AVFs, one important question remains unanswered: Do neointimal cells in maturation and failure have different origins or different paths of differentiation?

The limitations of this study include the single center setting and the exclusion of forearm AVFs. Human veins and fistulas may contain blood trapped in microvessels that can potentially contribute to small inflammatory clusters that were not analyzed in this study. We are aware of cell capture biases in scRNA-seq. To overcome this limitation, we presented with caution global cell proportions and only compared relative percentages within each major cluster. We also looked for further confirmation using standard histological techniques.

In conclusion, our study overcomes the limitations of bulk RNA-seq and provides an in-depth characterization of cellular populations in the human vein before and after arterialization. Our findings support the future testing of anti-inflammatory and anti-fibrotic agents to improve AVF maturation. We also uncover cell-specific and outcome-associated phenotypic changes that can advance the design of cell-targeted therapies. This dataset is a powerful tool to validate druggable targets in clinically relevant tissues, guiding future mechanistic studies in animal models and target selection for translation.

## DATA SHARING STATEMENT

Sequencing data were deposited in the Gene Expression Omnibus, accession number GSEXXXXXX (pending).

---

**Supplementary Table S1.**
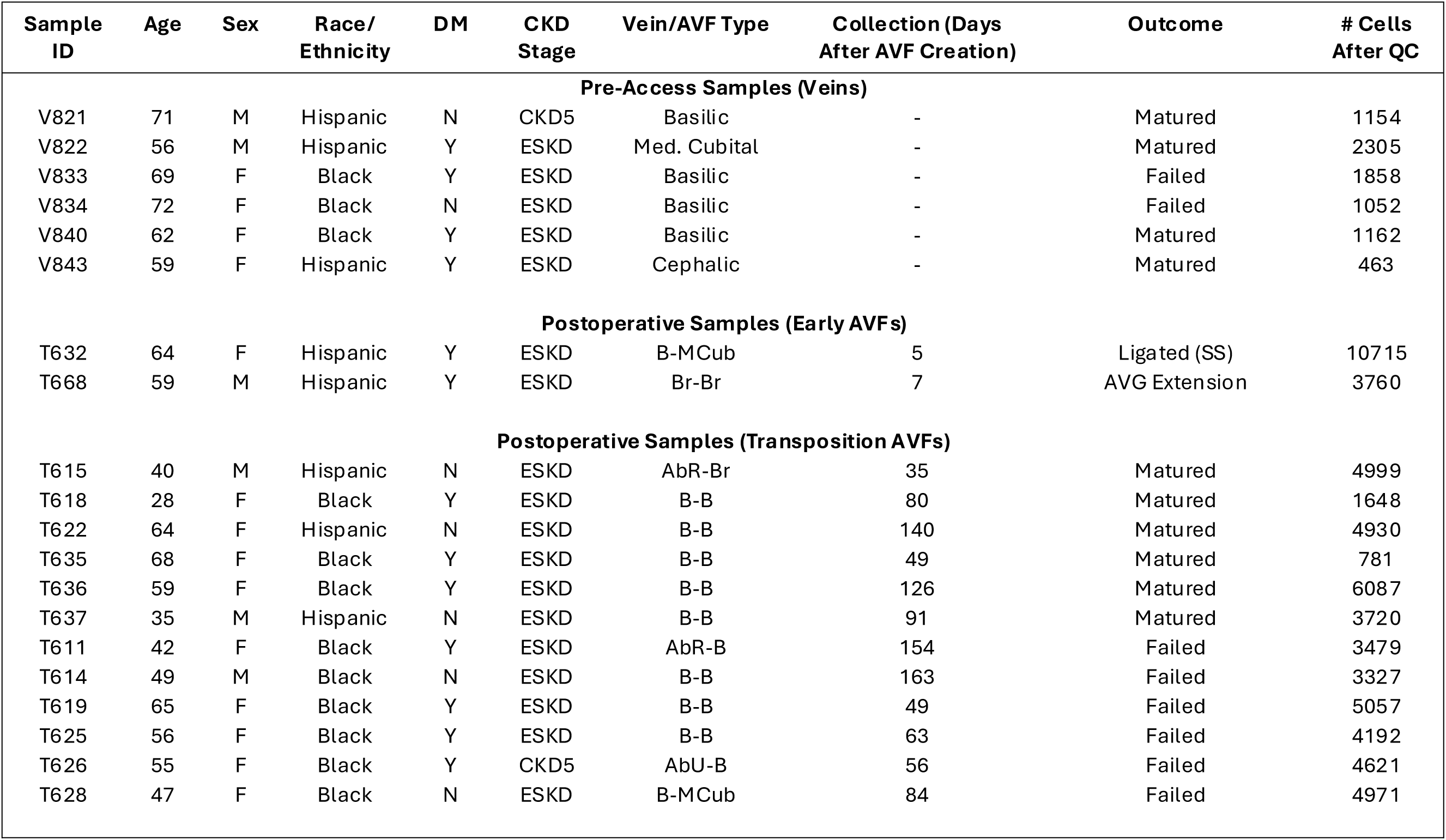
Characteristics of vein and AVF patients in the single-cell RNA sequencing cohort.

**Supplementary Table S2.** Patient characteristics in the paired vein-AVF validation cohort.

| Characteristics | All (n=23) | Matured (n=11) | Failed (n=12) |
| --- | --- | --- | --- |
| **Demographic** |  |  |  |
| Age (years, mean ± SD) | 61 ± 11 | 62 ± 12 | 61 ± 11 |
| Female (%) | 12 (52) | 5 (45) | 7 (58) |
| Race/ethnicity (%) |  |  |  |
| Black | 12 (52) | 4 (36) | 8 (67) |
| Hispanic | 6 (26) | 4 (36) | 2 (17) |
| White | 5 (22) | 3 (27) | 2 (17) |
| **Clinical history** |  |  |  |
| HTN (%) | 22 (96) | 10 (91) | 12 (100) |
| DM (%) | 15 (65) | 7 (64) | 8 (67) |
| CAD (%) | 6 (26) | 3 (27) | 3 (25) |
| CHF (%) | 5 (22) | 1 (9) | 4 (33) |
| CVA/TIA (%) | 5 (22) | 2 (18) | 3 (25) |
| ESKD (%) | 15 (65) | 6 (55) | 9 (75) |
| **Vascular access** |  |  |  |
| AVF type (%) |  |  |  |
| Brachiobasilic | 18 (78) | 9 (82) | 9 (75) |
| Aberrant radio-basilic | 2 (9) | 1 (9) | 1 (8) |
| Aberrant radio-cephalic | 1 (4) | 1 (9) | - |
| Brachiocephalic | 1 (4) | - | 1 (8) |
| Brachio-brachial | 1 (4) | - | 1 (8) |
| Time between vein and AVF collection (days, mean ± SD) | 83 ± 36 | 76 ± 29 | 89 ± 41 |
| Catheter use (%) | 15 (65) | 6 (55) | 9 (75) |
| Previous AVF (%) | 5 (22) | 2 (18) | 3 (25) |

**Table S3.** Cell proportions expressed as percentage of total cells in veins and second-stage AVFs.

|  |  |  |  |
| --- | --- | --- | --- |
| Cluster | Vein, n=6<br>(Ave ± SD) | AVF, n=12<br>(Ave ± SD) | P |
| EC | 24.8 ± 14.6 | 16.5 ± 9.6 | 0.17 |
| SMC | 4.4 ± 3.2 | 7.2 ± 8.3 | 0.49 |
| Myofibro | 7.1 ± 3.4 | 9.6 ± 4.7 | 0.47 |
| Fibroblast | 11.4 ± 5.0 | 21.6 ± 9.4 | 0.060 |
| Pericyte | 4.8 ± 4.0 | 4.8 ± 4.7 | 0.77 |
| Mono/Mac | 23.9 ± 15.8 | 19.1 ± 10.7 | 0.66 |
| B Cell | 0.3 ± 0.3 | 0.8 ± 0.9 | 0.37 |
| NK/T Cell | 16.3 ± 6.4 | 13.5 ± 9.0 | 0.43 |
| Mast Cell | 5.8 ± 1.6 | 2.1 ± 1.6 | 0.0031 |
| Neutrophil | 1.1 ± 1.9 | 4.8 ± 6.2 | 0.049 |

**Table S4.** Cell proportions expressed as percentage of total cells in early AVFs.

|  |  |
| --- | --- |
| Cluster | Early AVF, n=2<br>(Range) |
| EC | 10.2 – 12.1 |
| SMC | 2.6 – 3.0 |
| Myofibro | 11.3 – 12.6 |
| Fibroblast | 5.7 – 15.7 |
| Pericyte | 1.4 – 3.0 |
| Mono/Mac | 28.9 – 38.4 |
| B Cell | 0.7 – 1.0 |
| NK/T Cell | 13.6 – 17.2 |
| Mast Cell | 1.5 – 3.7 |
| Neutrophil | 0.7 – 3.2 |

**Table S5.** Relative proportions of mono/mac subtypes expressed as percentage of total cells in the cluster in veins and second-stage AVFs.

|  |  |  |  |
| --- | --- | --- | --- |
| Subcluster | Vein, n=6<br>(Ave ± SD) | AVF, n=12<br>(Ave ± SD) | P |
| Mac_1 | 52.7 ± 7.9 | 25.6 ± 12.4 | <0.001 |
| Mac_2 | 9.0 ± 5.5 | 12.1 ± 6.1 | 0.24 |
| Mac_3 | 25.0 ± 6.1 | 35.3 ± 15.9 | 0.20 |
| Mac_4 | 1.2 ± 1.5 | 7.8 ± 11.5 | 0.0064 |
| Mono | 1.5 ± 1.4 | 11.2 ± 7.2 | <0.0001 |
| DC | 10.6 ± 7.4 | 8.0 ± 4.5 | 0.45 |

**Table S6.**
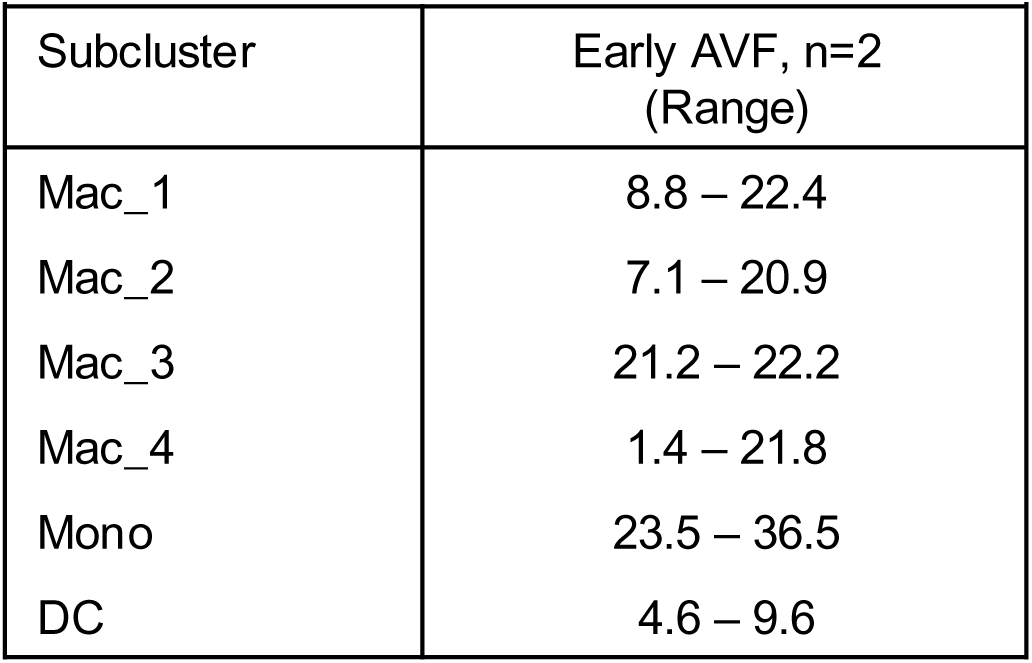
Proportions of mono/mac subtypes expressed as percentage of total cells in the cluster in early AVFs.

**Table S7.** Relative proportions of EC subtypes expressed as percentage of total cells in the cluster in veins and second-stage AVFs.

| Subcluster | Vein, n=6<br>(Ave ± SD) | AVF, n=12<br>(Ave ± SD) | P |
| --- | --- | --- | --- |
| EC_1 | 66.9 ± 21.5 | 63.6 ± 27.7 | 0.97 |
| EC_2 | 20.0 ± 9.6 | 4.5 ± 3.5 | 0.0033 |
| EC_3 | 7.1 ± 14.5 | 3.1 ± 4.7 | 0.43 |
| EC_4 | 2.4 ± 3.6 | 19.8 ± 20.6 | 0.0057 |
| EC_5 | 3.6 ± 2.4 | 9.0 ± 11.1 | 0.25 |

**Table S8.** Proportions of EC subtypes expressed as percentage of total cells in the cluster in early AVFs.

| Subcluster | Early AVF, n=2<br>(Range) |
| --- | --- |
| EC_1 | 19.6 – 64.2 |
| EC_2 | 6.2 – 6.7 |
| EC_3 | 4.6 – 5.1 |
| EC_4 | 21.6 – 27.6 |
| EC_5 | 3.3 – 41.0 |

**Table S9.**
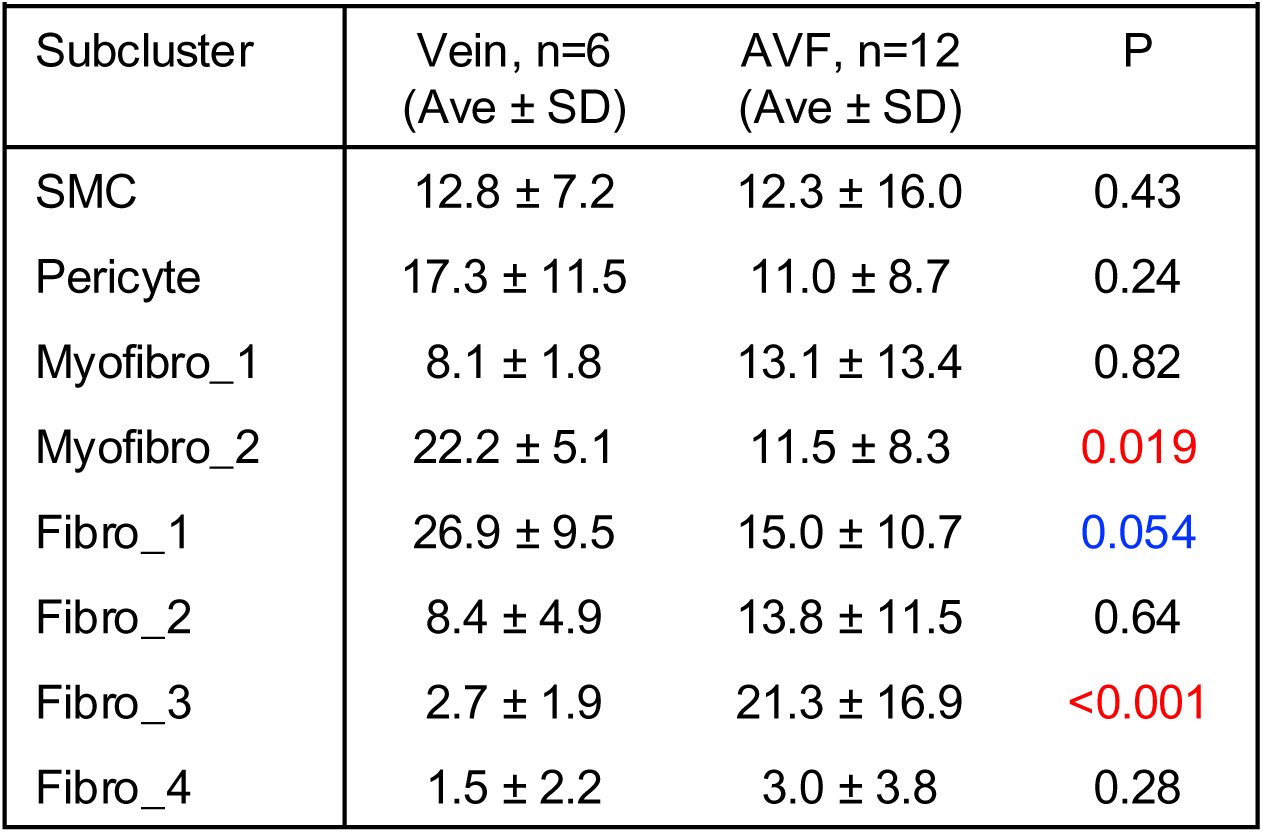
Relative proportions of mural cells and fibroblast subtypes expressed as percentage of total cells in the cluster in veins and second-stage AVFs.

**Table S10.** Proportions of mural cells and fibroblast subtypes expressed as % of total cells in the cluster in early AVFs.

| Subcluster | Early AVF, n=2<br>(Range) |
| --- | --- |
| SMC | 2.2 – 2.6 |
| Pericyte | 4.6 – 11.1 |
| Myofibro_1 | 14.5 – 16.8 |
| Myofibro_2 | 1.9 – 8.5 |
| Fibro_1 | 0.7 – 1.9 |
| Fibro_2 | 4.5 – 3.8 |
| Fibro_3 | 12.4 – 37.3 |
| Fibro_4 | 18.5 – 59.0 |

**Table S11.**
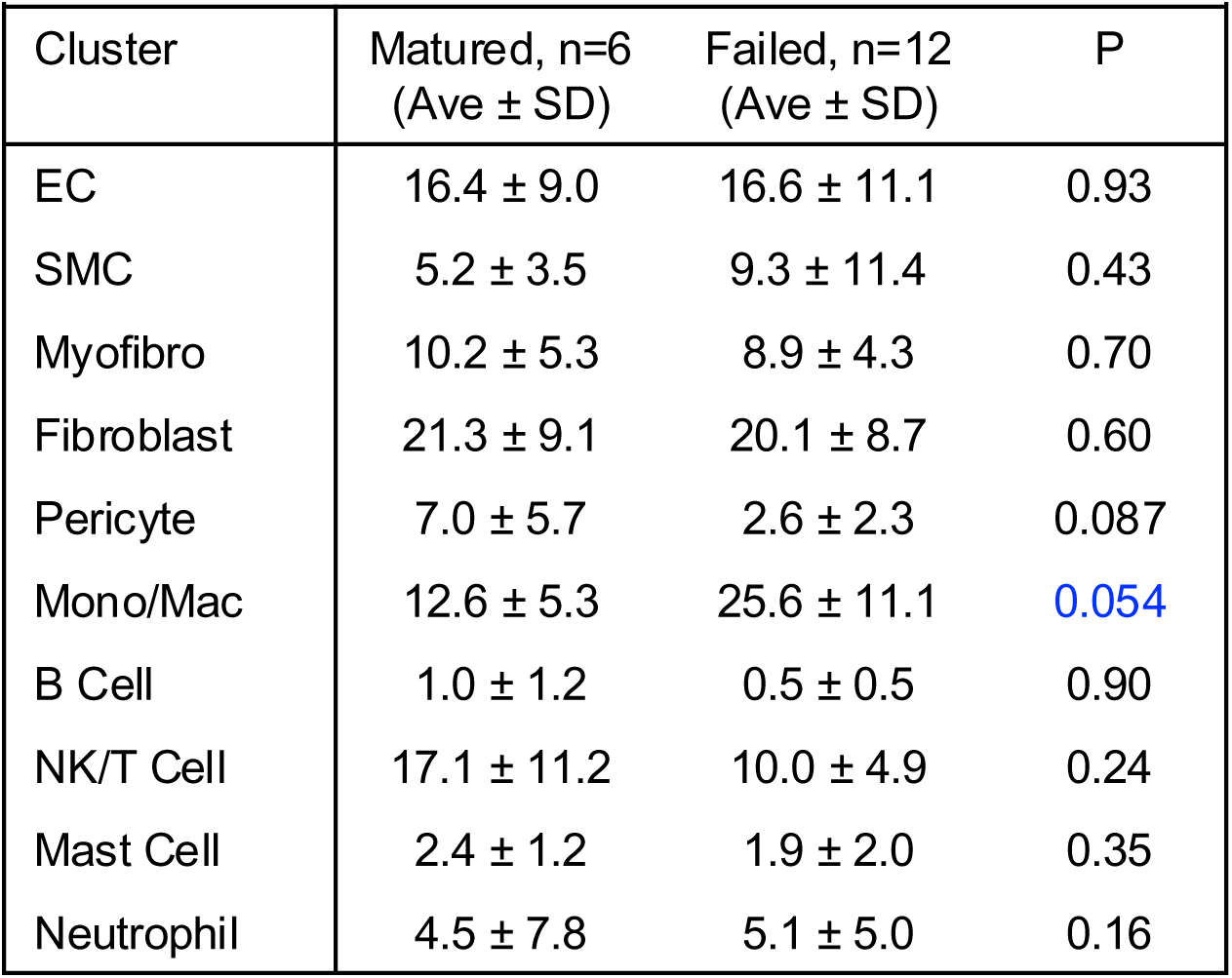
Cell proportions expressed as percentage of total cells in mature and failed AVFs.

| Cluster | Matured, n=6<br>(Ave ± SD) | Failed, n=12<br>(Ave ± SD) | P |
| --- | --- | --- | --- |
| EC | 16.4 ± 9.0 | 16.6 ± 11.1 | 0.93 |
| SMC | 5.2 ± 3.5 | 9.3 ± 11.4 | 0.43 |
| Myofibro | 10.2 ± 5.3 | 8.9 ± 4.3 | 0.70 |
| Fibroblast | 21.3 ± 9.1 | 20.1 ± 8.7 | 0.60 |
| Pericyte | 7.0 ± 5.7 | 2.6 ± 2.3 | 0.087 |
| Mono/Mac | 12.6 ± 5.3 | 25.6 ± 11.1 | 0.054 |
| B Cell | 1.0 ± 1.2 | 0.5 ± 0.5 | 0.90 |
| NK/T Cell | 17.1 ± 11.2 | 10.0 ± 4.9 | 0.24 |
| Mast Cell | 2.4 ± 1.2 | 1.9 ± 2.0 | 0.35 |
| Neutrophil | 4.5 ± 7.8 | 5.1 ± 5.0 | 0.16 |

**Table S12.** Relative proportions of mono/mac subtypes expressed as percentage of total cells in the cluster in mature and failed AVFs.

| Subcluster | Vein, n=6<br>(Ave ± SD) | AVF, n=12<br>(Ave ± SD) | P |
| --- | --- | --- | --- |
| Mac_1 | 25.6 ± 14.1 | 25.6 ± 11.9 | 0.87 |
| Mac_2 | 12.8 ± 5.1 | 11.4 ± 7.4 | 0.59 |
| Mac_3 | 30.4 ± 8.8 | 40.3 ± 20.5 | 0.35 |
| Mac_4 | 10.3 ± 16.1 | 5.3 ± 3.7 | 0.89 |
| Mono | 10.7 ± 6.3 | 11.8 ± 8.5 | 0.99 |
| DC | 10.3 ± 5.4 | 5.7 ± 1.5 | 0.081 |

**Table S13.** Relative proportions of EC subtypes expressed as percentage of total cells in the cluster in mature and failed AVFs.

| Subcluster | Vein, n=6<br>(Ave ± SD) | AVF, n=12<br>(Ave ± SD) | P |
| --- | --- | --- | --- |
| EC_1 | 73.3 ± 26.2 | 54.0 ± 27.8 | 0.17 |
| EC_2 | 3.2 ± 3.0 | 5.7 ± 3.7 | 0.37 |
| EC_3 | 2.2 ± 2.5 | 4.0 ± 6.3 | 0.89 |
| EC_4 | 9.5 ± 12.1 | 30.2 ± 23.1 | 0.037 |
| EC_5 | 11.8 ± 15.7 | 6.2 ± 2.0 | 0.98 |

**Table S14.** Relative proportions of mural cells and fibroblast subtypes expressed as percentage of total cells in the cluster in mature and failed AVFs.

| Subcluster | Vein, n=6<br>(Ave ± SD) | AVF, n=12<br>(Ave ± SD) | P |
| --- | --- | --- | --- |
| SMC | 7.8 ± 4.9 | 16.7 ± 22.2 | 0.55 |
| Pericyte | 15.5 ± 9.9 | 6.5 ± 4.7 | 0.20 |
| Myofibro_1 | 12.1 ± 16.2 | 14.1 ± 11.3 | 0.66 |
| Myofibro_2 | 13.4 ± 9.0 | 9.7 ± 7.9 | 0.48 |
| Fibro_1 | 15.5 ± 9.6 | 14.5 ± 12.5 | 0.75 |
| Fibro_2 | 14.4 ± 9.3 | 13.3 ± 14.3 | 0.61 |
| Fibro_3 | 20.0 ± 18.5 | 22.6 ± 16.8 | 0.99 |
| Fibro_4 | 3.8 ± 5.2 | 2.2 ± 2.0 | 0.79 |

**Supplementary Figure S1.**
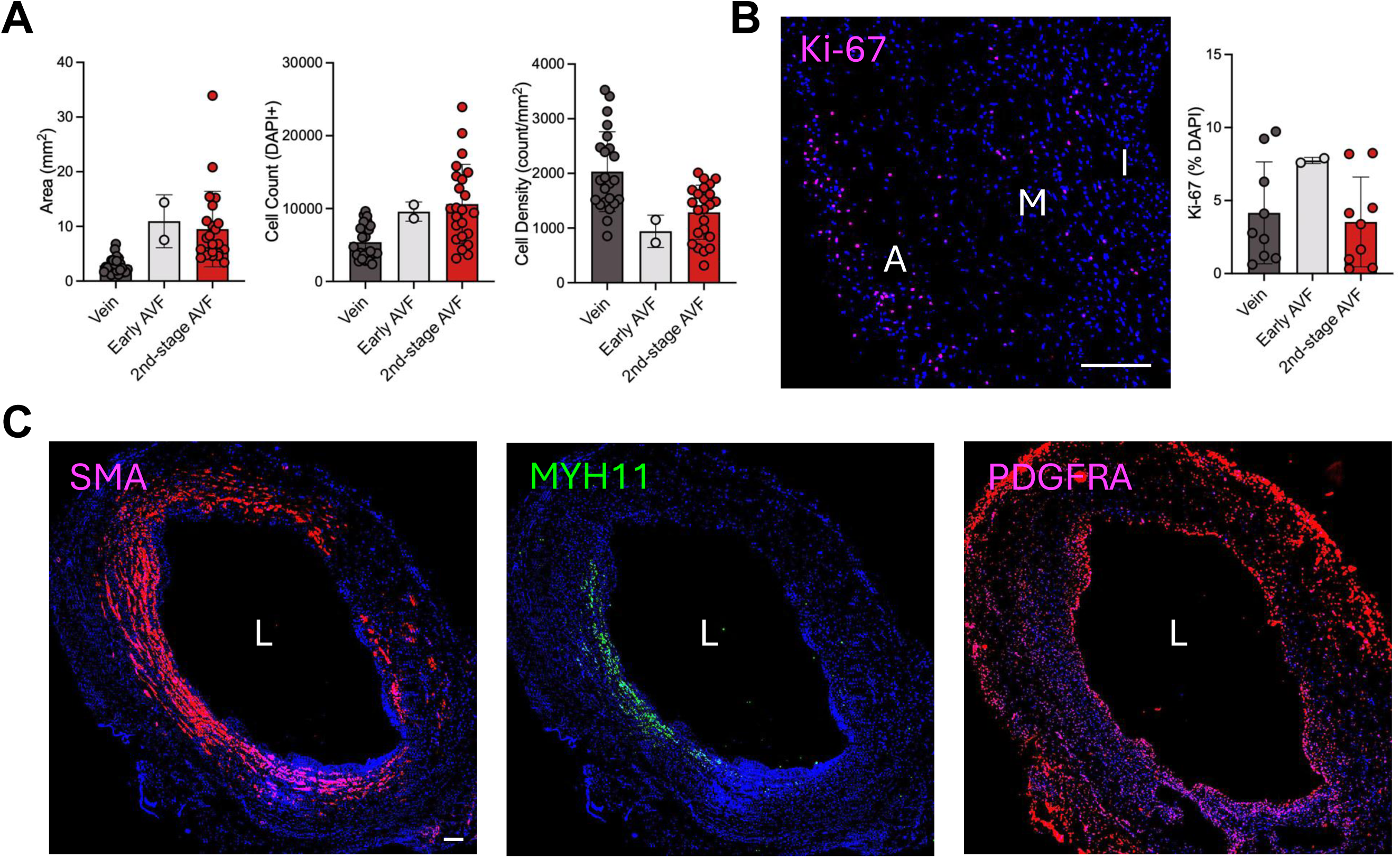
Wall morphometry and composition in one-week AVFs. **A)** Wall morphometry and cell density in pre-access veins, early AVFs, and second-stage AVFs. **B)** Ki-67 staining in early AVFs and quantitative comparison among tissue types. **C)** Distribution of contractile SMCs (MYH11+), synthetic SMCs or myofibroblasts (SMA+ MYH11-), and fibroblasts (PDGFRA+) in early AVFs. L, lumen. Scale bars = 200 um.

**Supplementary Figure S2.**
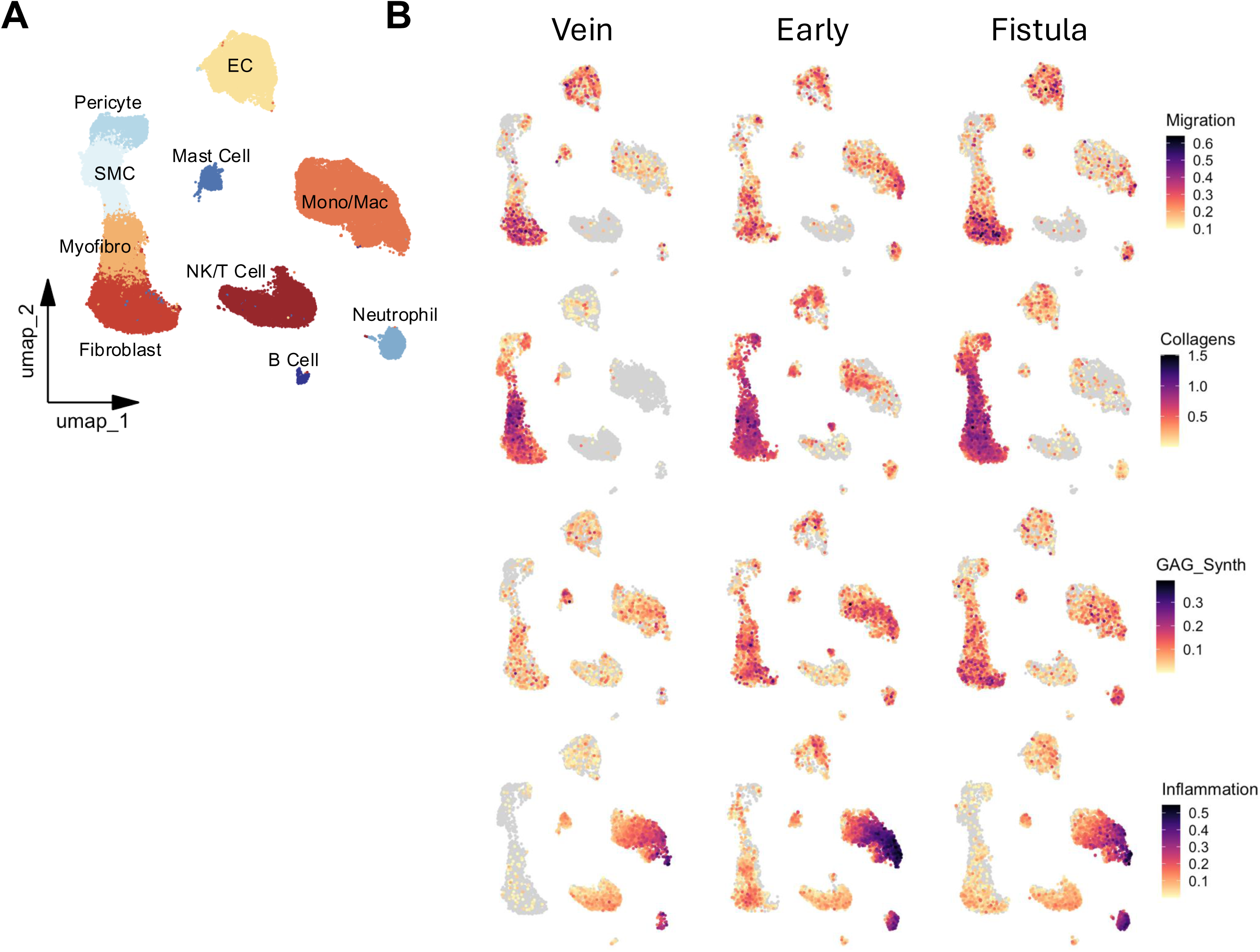
Functional profiles of main vascular clusters by scRNA-seq. **A)** UMAP of 70,281 cells from pre-access veins and AVFs. **B)** Functional profiling of single cell clusters in veins, early AVFs, and second-stage AVFs according to gene signature scores. 8,000 cells are projected per tissue type.

**Supplementary Figure S3.**
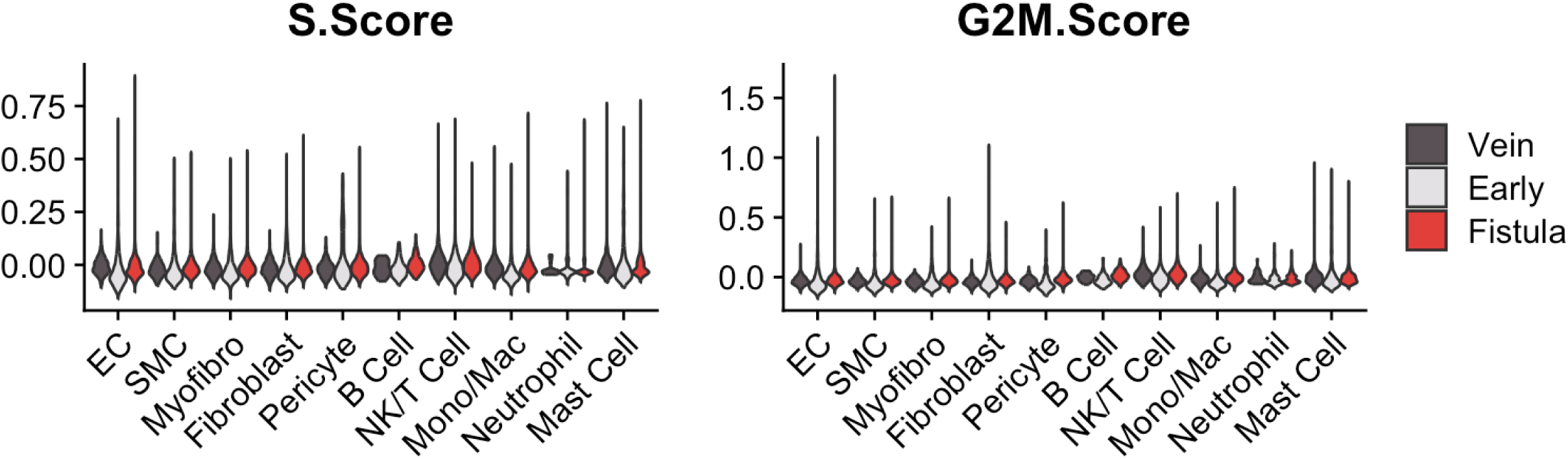
scRNA-seq predicts low proliferative activity in fistulas. S-phase (DNA synthesis) and G2/M-phase (cell division) scores in pre-access veins, early AVFs, and second-stage AVFs.

**Supplementary Figure S4.**
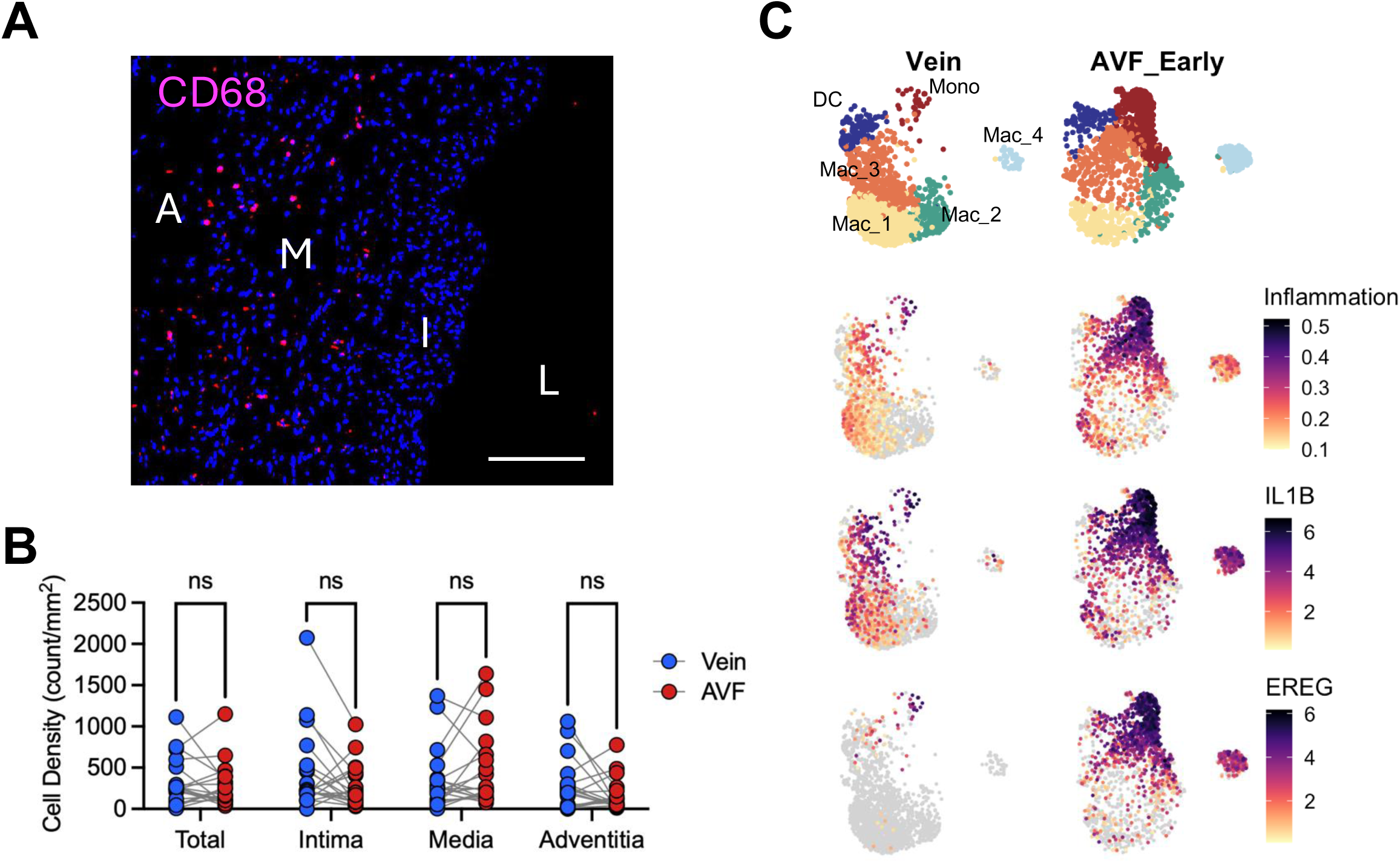
Macrophages in early AVFs. **A)** Infiltration of macrophages in early fistulas. L, lumen; I, intima; M, media; A, adventitia. Scale bar = 200 um. **B)** Macrophage density in vein and AVF pairs. **C)** Focused UMAPs of monocytic phenotypes in pre-access veins and early AVFs. 2,000 cells are projected per tissue type. Inflammatory monocytes appear after fistula creation.

**Supplementary Figure S5.**
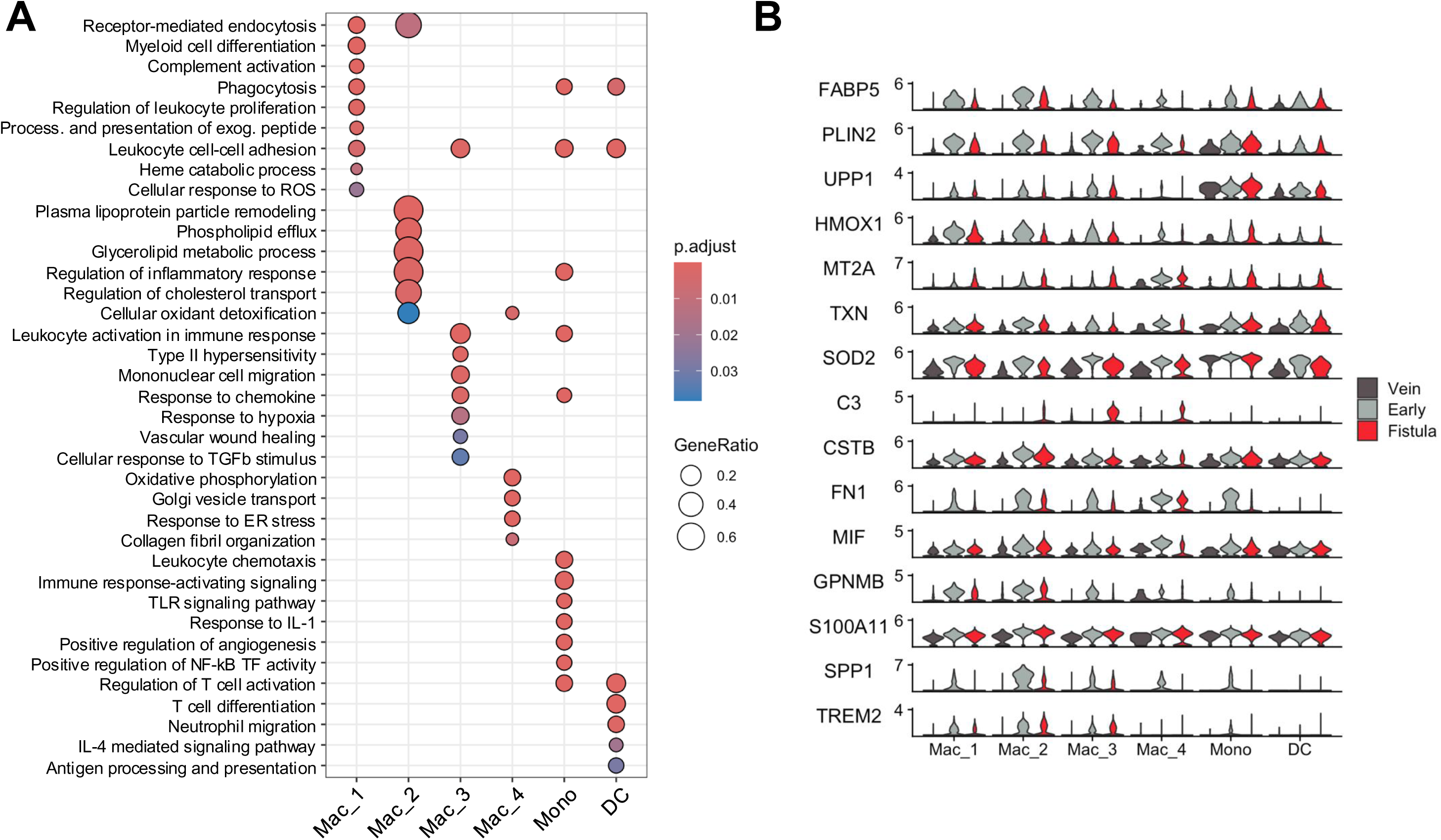
Functional characteristics of mono/mac phenotypes. **A)** Gene ontology (GO) over-representation analysis of biological processes enriched in mono/mac phenotypes. **B)** Upregulation of genes involved in inflammation, oxidative stress responses, and vascular healing across all phenotypes after AVF creation.

**Supplementary Figure S6.**
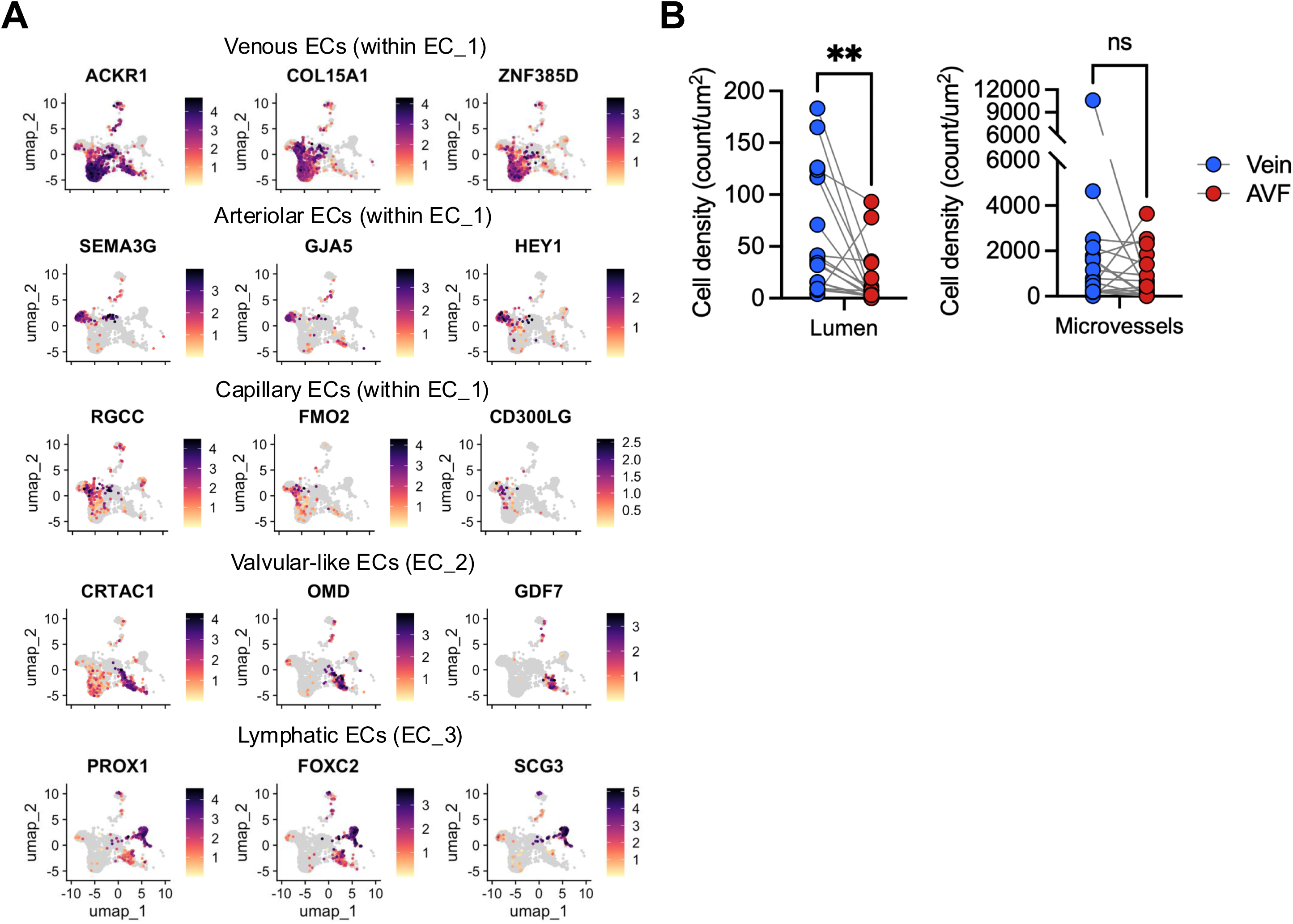
Endothelial phenotypes in pre-access veins. **A)** Expression markers defining the main types of ECs in pre-access veins. **B)** Quantification of ACKR3+ cells in vein and AVF pairs. **p<0.01

**Supplementary Figure S7.**
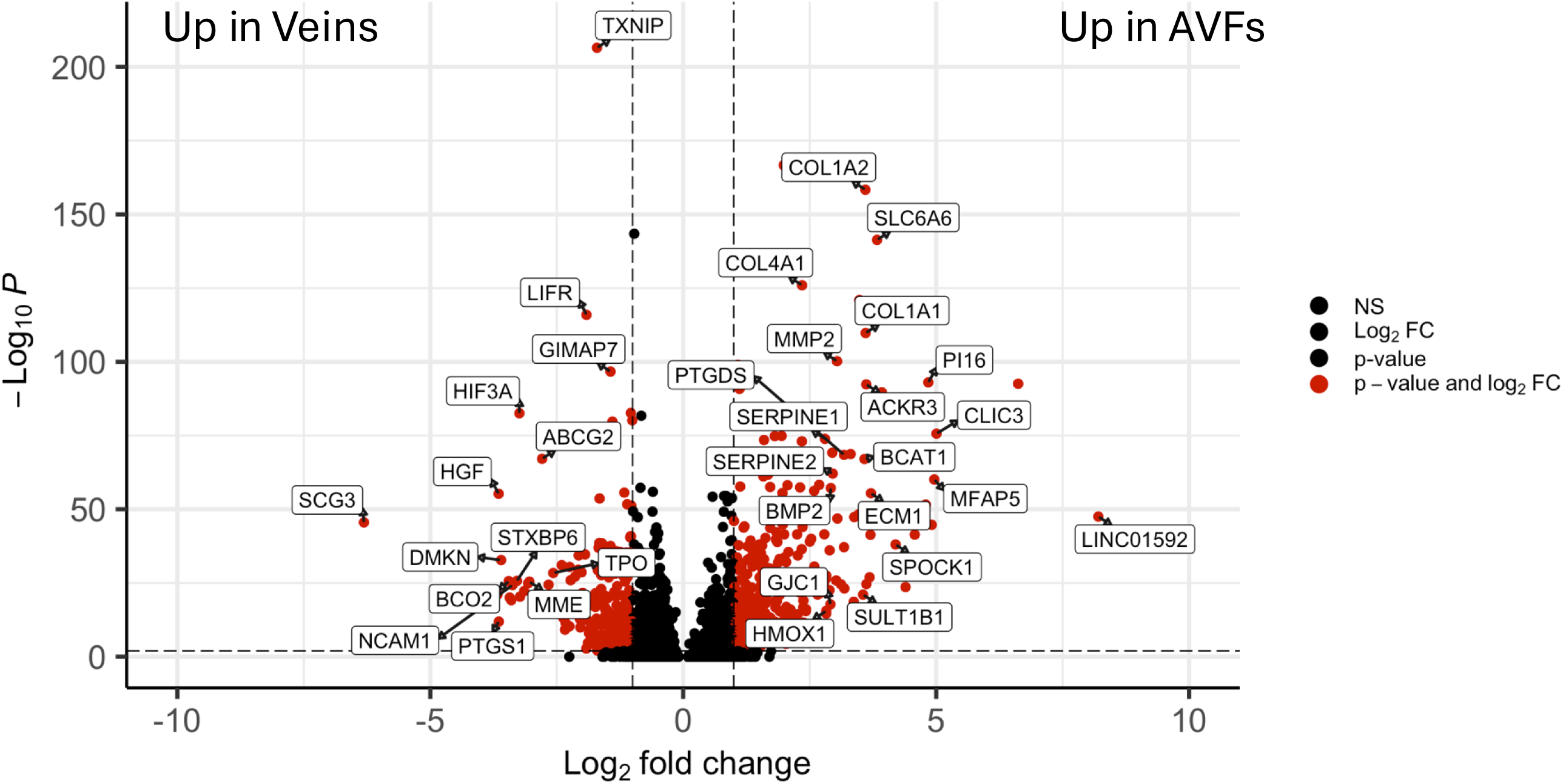
Differentially expressed genes in ECs from pre-access veins and AVFs.

**Supplementary Figure S8.**
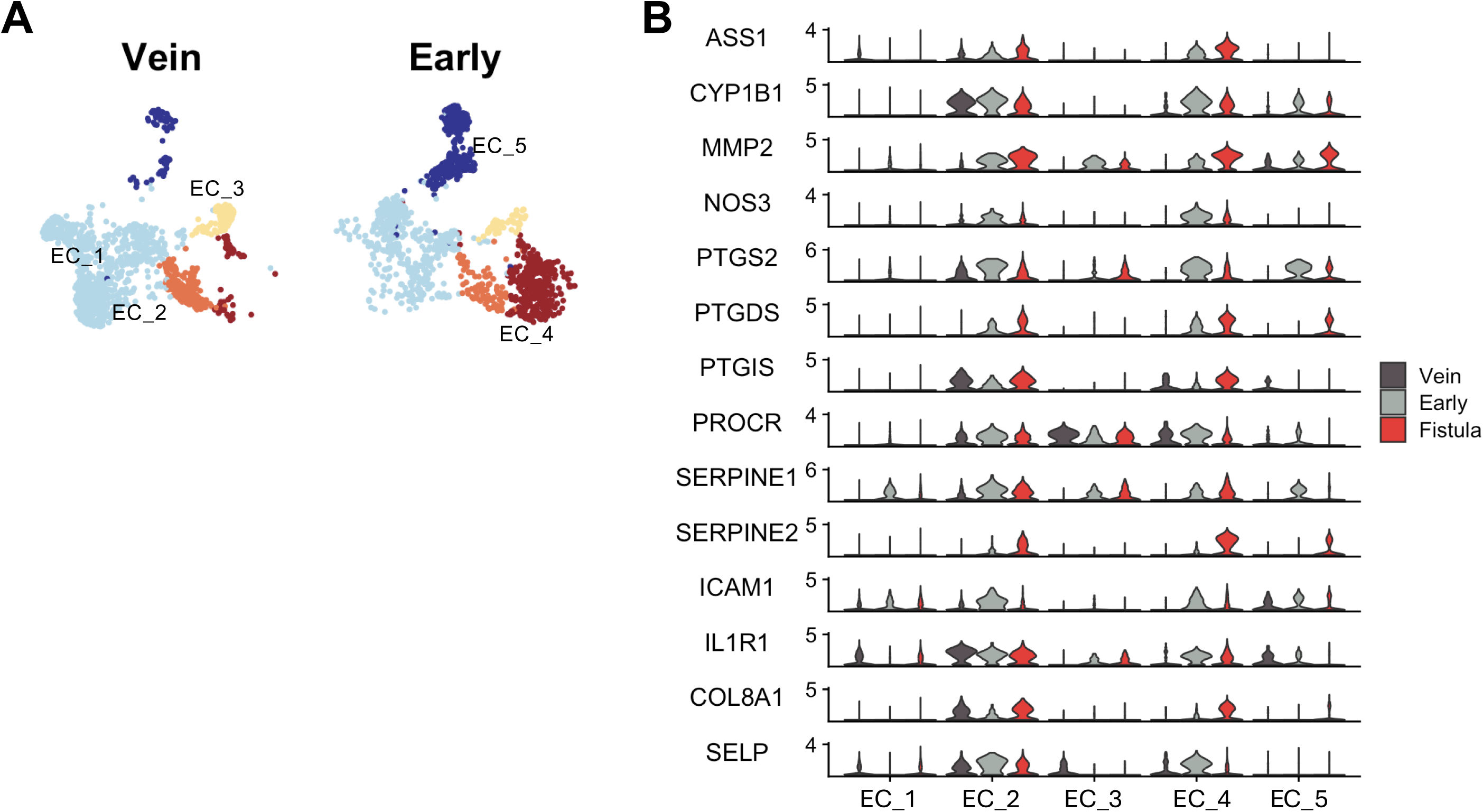
Early adaptations in ECs after fistula creation. **A)** Focused UMAPs of EC phenotypes in pre-access veins and early AVFs. 1,500 cells are projected per tissue type. **B)** Expression changes in selected shear stress sensing, hemostatic, and inflammatory genes in ECs from veins, early AVFs, and second-stage AVFs.

**Supplementary Figure S9.**
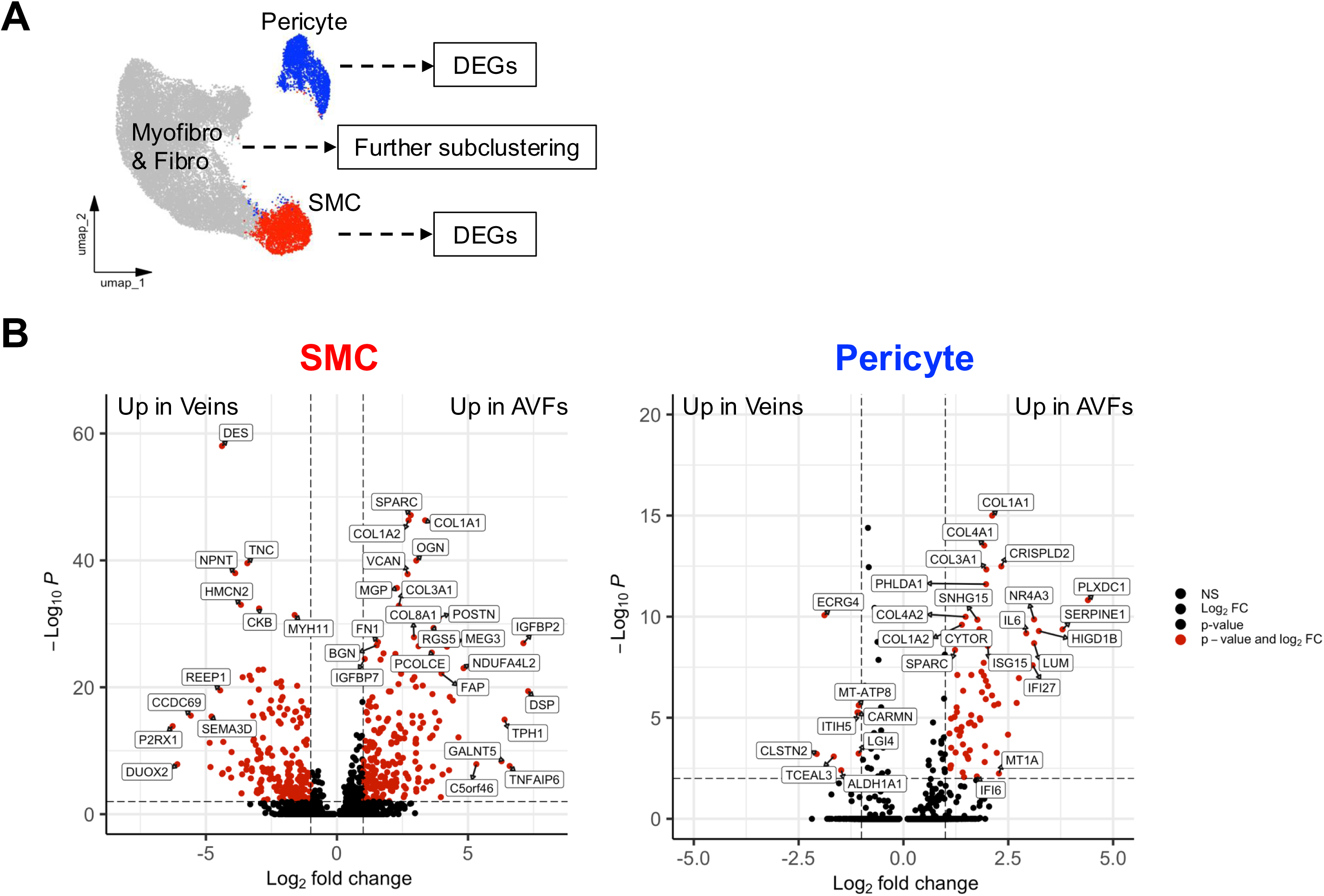
Mural cells in veins and AVFs. **A)** Re-clustering strategy for differential expression analyses of mural cells and further subclustering of myofibroblasts and fibroblasts. **B)** Differentially expressed genes in SMCs and pericytes from pre-access veins and AVFs.

**Supplementary Figure S10.**
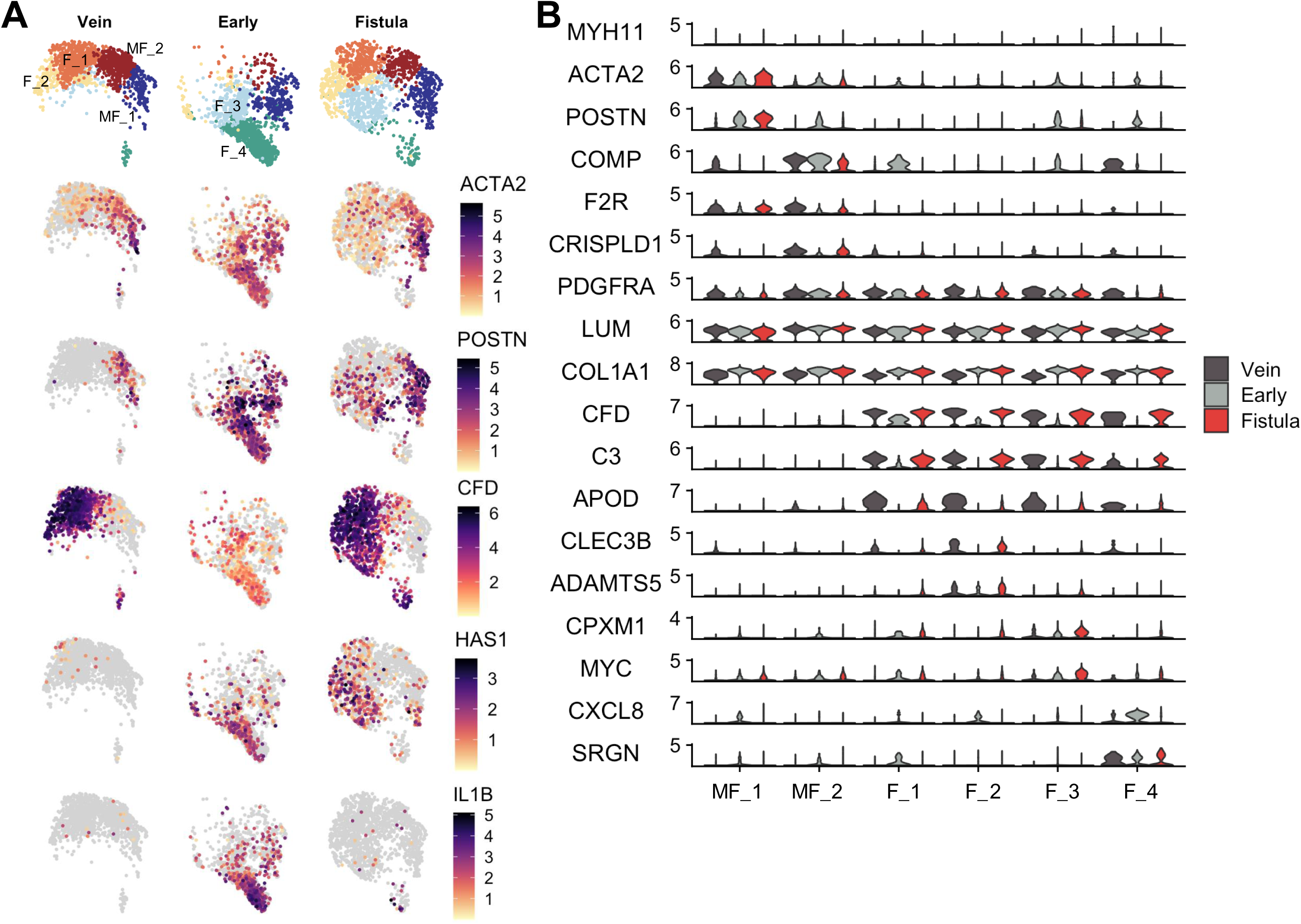
Early activation of healing programs in myofibroblasts (MF) and fibroblasts (F) after fistula creation. **A-B)** Expression markers identifying myofibroblasts and fibroblasts, and early changes after AVF creation. UMAPs show 1,500 cells per tissue type.

**Supplementary Figure S11.**
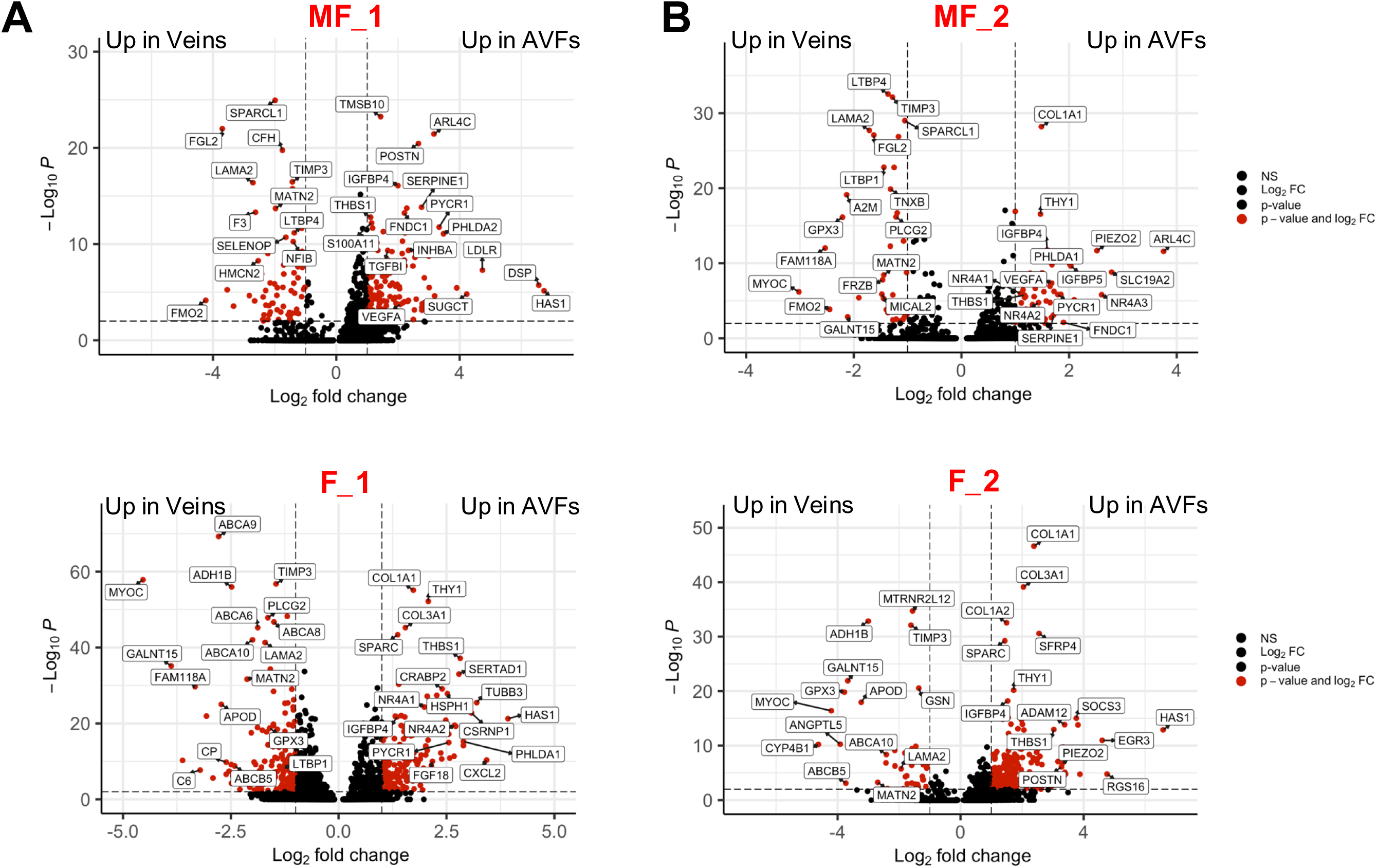
Differential gene expression analysis of myofibroblast and fibroblast subpopulations present in both pre-access veins and AVFs.

**Supplementary Figure S12.**
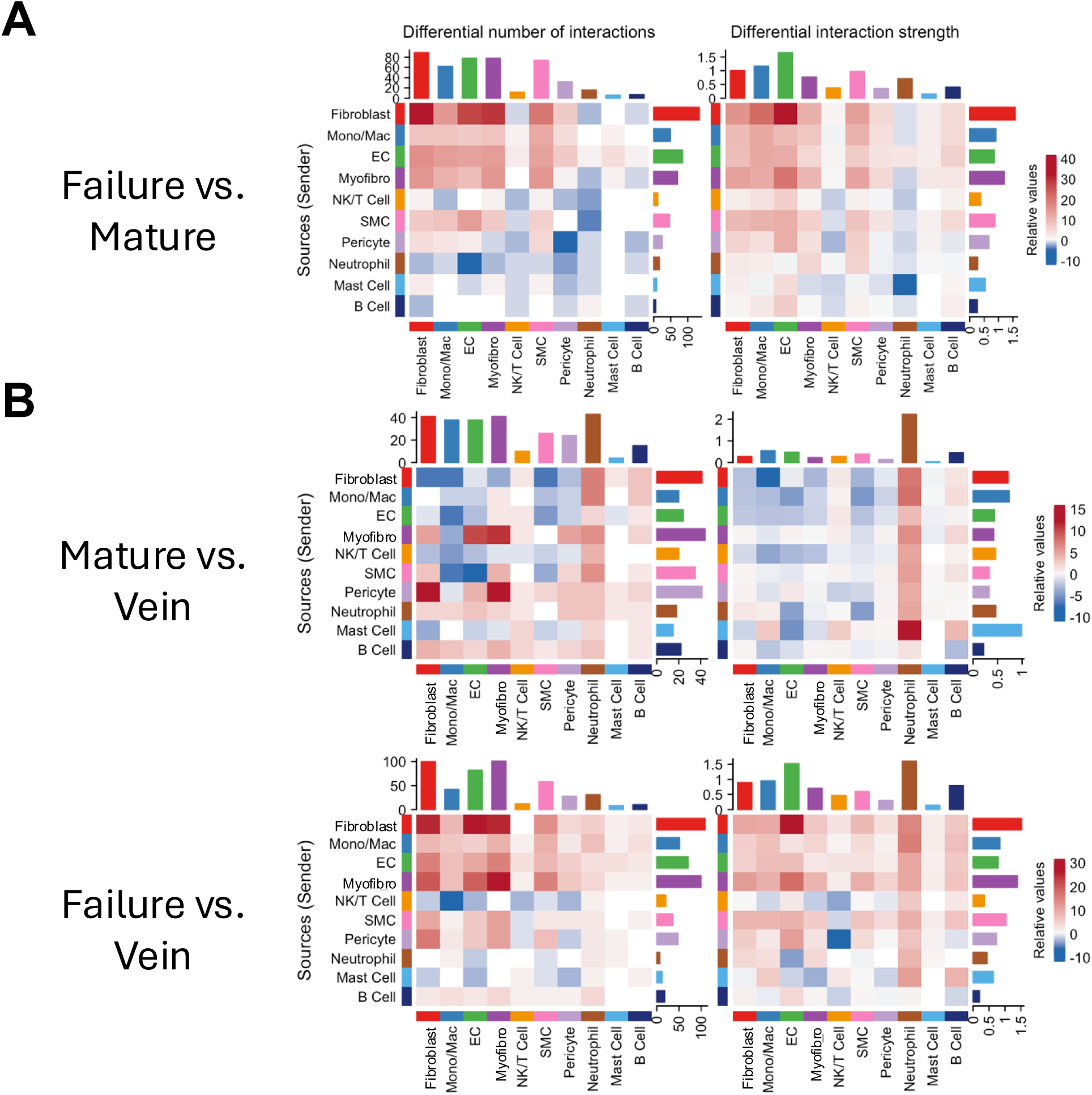
Comparative cell-to-cell communication analyses using CellChat. **A)** Communications among fibroblasts, mono/macs, ECs, and myofibroblasts are enhanced in failed vs. mature AVFs. **B)** Enhanced communications in failed AVFs compared to veins but in mature AVFs relative to veins.

**Supplementary Figure S13.**
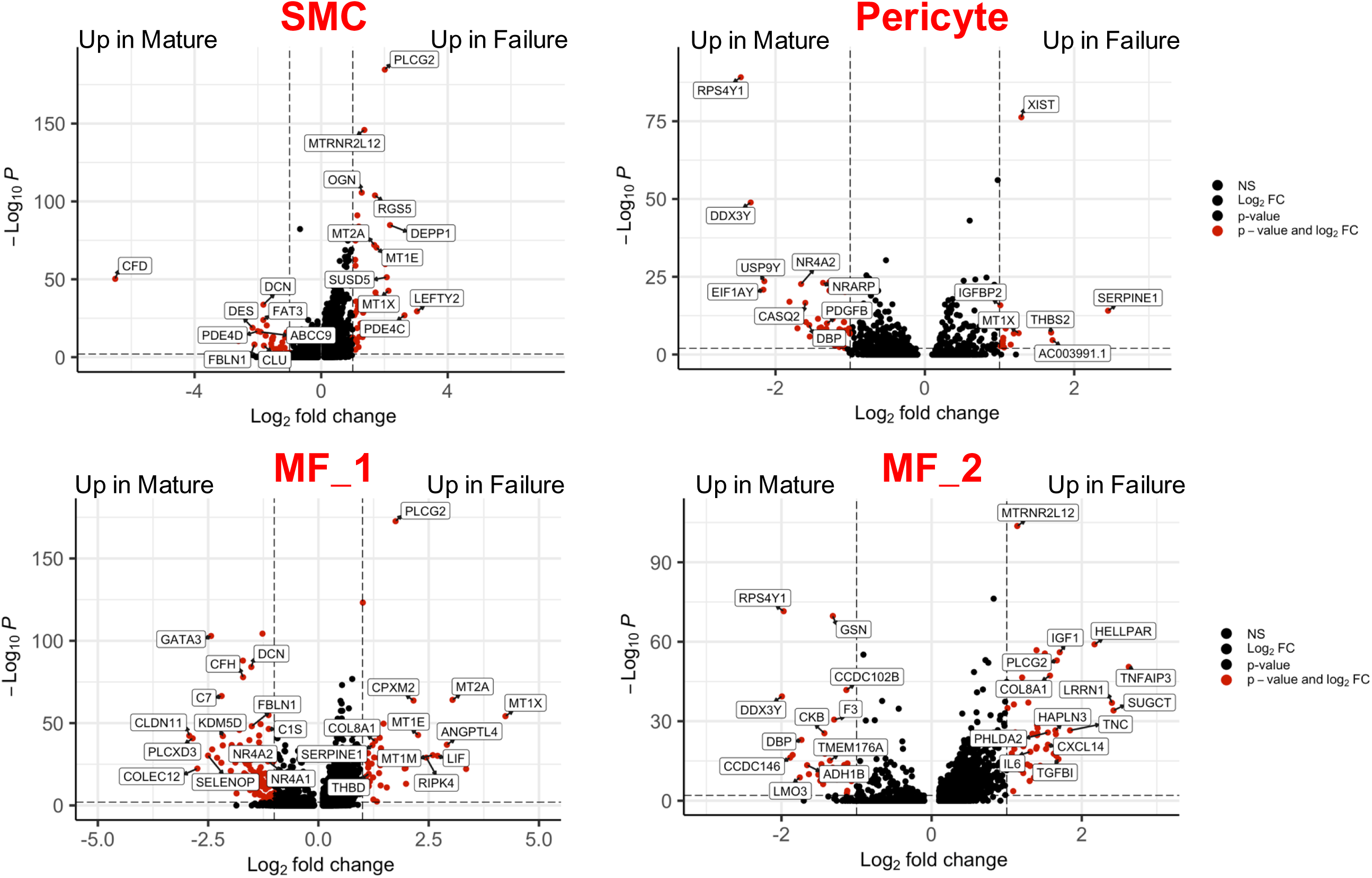
Differential gene expression analysis of mural cells and myofibroblasts from AVFs that matured and failed.

**Supplementary Figure S14.**
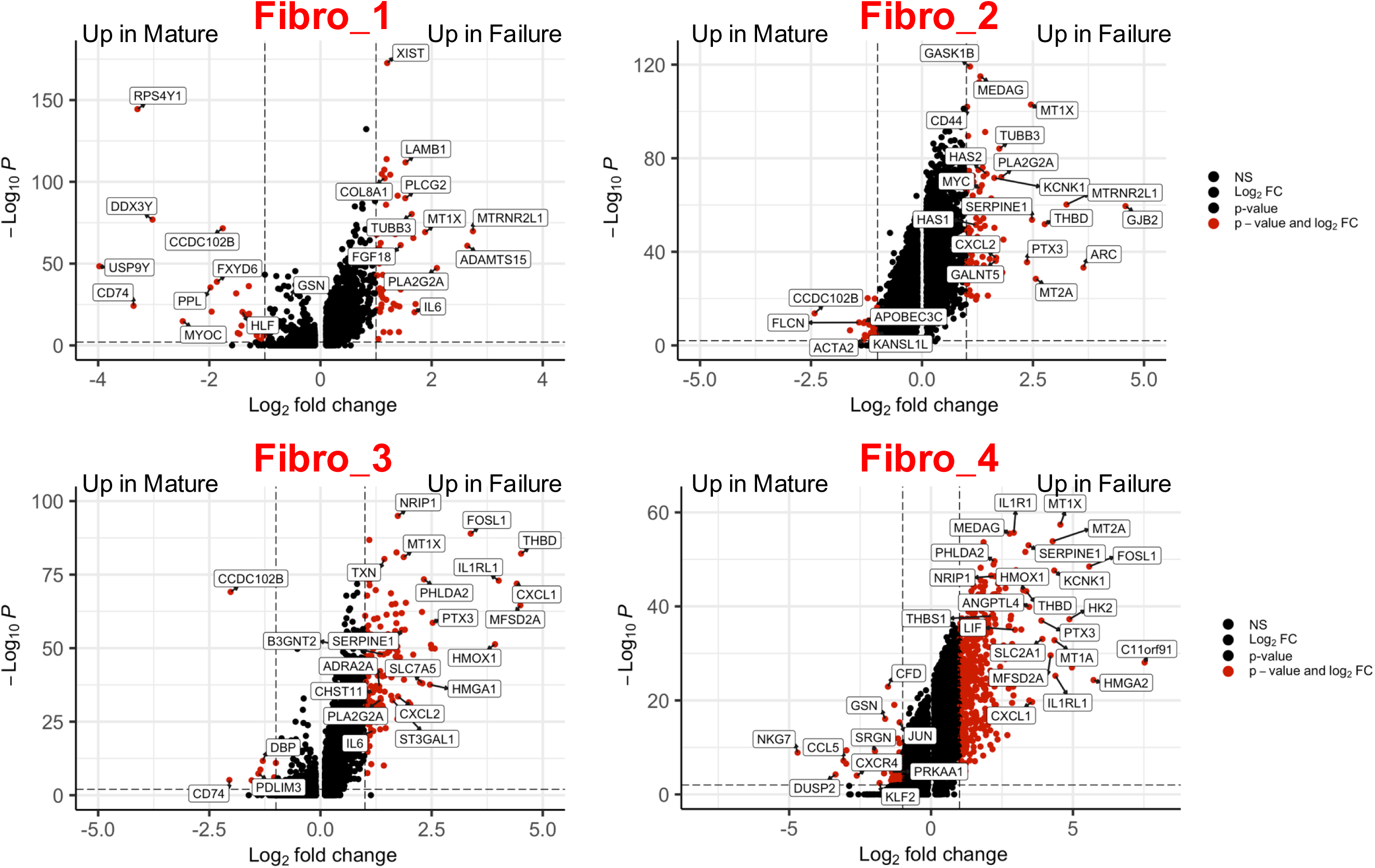
Differential gene expression analysis of myofibroblast phenotypes from AVFs that matured and failed.

## SUPPLEMENTARY METHODS

### Study Subjects and Sample Collection

For single-cell RNA sequencing (scRNA-seq), we enrolled 20 participants ≥21 years of age, with CKD5 or ESKD, and planned AVF creation in two stages at Jackson Memorial Hospital (JMH) or the University of Miami Hospital (UMH) from April 2022 to February 2024 (**Supplementary Table S1**). From these patients we obtained: 1) six pre-access veins collected at the time of access creation, 2) 12 juxta-anastomotic AVF biopsies (six matured, six failed) collected 91 ± 44 days after creation at the time of transposition, and 3) two rare early AVFs resected within 7 days of fistula creation due to high flow steal syndrome or a planned arteriovenous graft (AVG) extension. All samples were full cross-sections 5-10 mm in length. Vessels were collected in cold DMEM/F12 (Gibco, Thermo Fisher Scientific, Waltham, MA) supplemented with 10% FBS (Sigma-Aldrich, St. Louis, MO), 1 mM sodium pyruvate, 100 U/mL penicillin, 100 ug/mL streptomycin, and 50 ug/mL gentamicin (all Gibco).

For immunofluorescence (IF) analyses, we included longitudinal vein and second-stage AVF samples from 23 additional individuals (11 matured, 12 failed) randomly selected from the University of Miami Vascular Access biorepository (**Supplementary Table S2**). These samples were collected in RNA*later* (QIAGEN, Germantown, MD) and stored at −80°C. A 1-2 mm cross-section was fixed in 10% neutral formalin (Sigma-Aldrich, St. Louis, MO) for paraffin embedding and sectioning.

Anatomic maturation failure was defined as a fistula whose transected length did not allow a standard transposition because of stenosis and required a short transposition, AVG extension, or ligation.^1,2^ None of the AVFs underwent endovascular or surgical interventions to assist maturation. The study was performed according to the ethical principles of the Declaration of Helsinki and regulatory requirements at JMH and UMH. The ethics committee and Institutional Review Board at the University of Miami approved the study.

### Tissue Dissociation

To generate single-cell suspensions for sequencing or primary cell isolation, vessels were cut longitudinally with a sterile scalpel and then into 1-2 mm square sections. Samples were enzymatically digested for 90 minutes at 37°C with shaking using a combination of 3 mg/mL collagenase type II, 0.25 mg/mL soybean trypsin inhibitor, 0.2 mg/mL elastase, 1.4 U/mL Dispase, 60 U/mL DNAse I (all Worthington Biochemical, Lakewood, NJ), 0.15 mg/mL collagenase type XI, 0.25 mg/mL hyaluronidase type I, and 2.38 mg/mL HEPES (all Sigma-Aldrich) in HBSS with Ca^2+^ and Mg^2+^ (Thermo Fisher). After 90 minutes, an equal volume of Accumax (Sigma-Aldrich) was added to the dissociation mix and incubated for an additional 5 minutes. Single cells were filtered through a 40 µM strainer, washed with HBSS twice, and incubated with red blood cell lysis buffer (BioLegend, San Diego, CA) for 5-10 min. Cells were washed once again with HBSS and resuspended in 0.1% BSA in PBS (Gibco) for counting and RNA sequencing, or in the corresponding media for primary cell culture.

### Single-Cell RNA Sequencing and Alignment

Preparation of single-cell RNA libraries and sequencing were performed in the Center for Genome Technology at the University of Miami John P. Hussman Institute for Human Genomics. Single cell suspensions were counted using both the Cellometer K2 Fluorescent Viability Cell Counter (Nexcelom) and a hemocytometer. Samples with >80% viability were run using the Chromium Single Cell 3′ Library & Gel Bead Kit v3 (10X Genomics). The manufacturer’s protocol was used with a target capture of 10,000 cells. Each sample was processed on an independent Chromium Single Cell A Chip (10X Genomics) and subsequently run on a thermocycler (Eppendorf). Sequencing libraries were evaluated for quality on the Agilent Tape Station (Agilent Technologies, Palo Alto, CA), quantified using a Qubit 2.0 Fluorometer (Invitrogen, Carlsbad, CA), and qPCR before sequencing on the Illumina NovaSeq 6000. FASTQ files were generated with Cell Ranger’s mkfastq pipeline (version 6.0.2). The Cell Ranger’s count pipeline (version 6.0.2) was used to generate raw gene-barcode matrices from alignment to the 10X Genomics pre-built Cell Ranger human reference package (version 2020-A), from GRCh38 Ensembl build 98/GENCODE v32 gene annotations.

### Bioinformatic Quality Control

Data were processed in R (4.2.2) using Seurat v4.^3^ Briefly, cells with >15% mitochondrial content, >50% ribosomal content, <200 genes (nFeature_RNA), low complexity (log_10_GenesPerUMI<0.80), or predicted as doublets according to DoubletFinder^4^ or gene count (nFeature_RNA>8000) were filtered out from downstream analyses. Data were normalized using the “LogNormalize” method and using a scale factor of 10,000. Using Seurat’s Scale.Data() function and “vars.to.regress” option, cell cycle, percent mitochondrial genes, and number of UMIs were used to regress out unwanted sources of variation. Unless noted otherwise, all bioinformatic packages were used as detailed in their respective vignettes with no major modifications to the R/Python Code.

### Cell Clustering and Functional Annotations

We merged all 20 vessels from our study, as well as four basilic/cephalic veins and three brachial arteries from non-CKD donors (GEO accession numbers GSE250469 and GSE266682)^5,6^ in Seurat. After applying the functions *NormalizeData*, *FindVariableFeatures*, *ScaleData*, and *RunPCA,* we removed batch effects and generated an integrated map using Harmony.^7^ Overall clustering was performed with *FindNeighboors* and the first 30 principal components, followed by *Findclusters* and *RunUMAP* at a 0.5 resolution. We manually annotated the main clusters according to canonical cell markers.^5,6^ From this general Seurat object, we then subset the 20 samples from this study for graphical representations and bioinformatic analyses of individual clusters.

Functional scores were added to the integrated Seurat object using the function *AddModuleScore* in Seurat 4.0. Scores were based on curated gene signature modules from the Molecular Signatures Database (Human MSigDB v2023.1.Hs)^8,9^, including the HALLMARK gene sets of Angiogenesis, Inflammatory Response, and Coagulation; the REACTOME module R-HSA-202733; and the GOBP modules GO:0051894, GO:0007263, GO:0034405, GO:0007160, and GO:0042060 R-HSA-9707564, R-HSA-1234174, and R-HSA-3299685. Additional scores were generated using the collagens and proteoglycans gene sets from the Matrisome Database.^10^ Cell proportions were compared using the propeller method.^11^

### Differential Gene Expression Analyses and Bioinformatic Inferences

Differential gene expression analyses of individual subclusters were performed using the *FindMarkers* function after random downsampling to include the same number of cells per tissue type in AVF vs. vein comparisons, or the same number of cells per individual in analyses in failed vs. mature AVFs. Ligand-receptor interactomes were analyzed using CellChat v2.^12^ We imported the clustering metadata and generated a CellChat object with all subclusters following the package vignette. Overrepresented interactions were calculated using *identifyOverExpressedInteractions* and only those present in at least 10 cells per cluster were retained for downstream analysis. Comparisons of different datasets was performed on a merged CellChat object using *compareInteractions*.

### Primary Cell Isolation and Culture

Primary AVF fibroblasts were isolated by enzymatic digestion of vessels as above, followed by CD45-/CD31-/CD90+ cell sorting using CD45, CD31, and CD90 microbeads (Miltenyi Biotec, North Rhine-Westphalia, Germany). Fibroblasts were cultured in CytoSoft flasks or plates with low rigidity (0.2 kPa; Advanced Biomatrix, Carlsbad, CA) using Fibroblast Growth Medium 2 (PromoCell, Heidelberg, Germany) to avoid spontaneous phenotypic activation. Cells were stimulated with TGF-β1 (R&D Systems, Minneapolis, MN) or IL-1β (PeproTech, Cranbury, NJ) for 24 hours, each at a final concentration of 10 ng/mL. For inhibitory conditions, cells were pre-incubated for 1 hour with an NF-kB inhibitor (10 uM; Cayman Chemical Co., Ann Arbor, MI; cat. #BMS345541) or a TGFβRI inhibitor (1 uM; Tocris Bioscience, cat. #SB525334) prior to cytokine treatment. Cell passages 5 to 7 were used in all experiments.

Human umbilical vein endothelial cells (HUVEC) were purchased from Lonza (Walkersville, MD) and cultured in EGM-2 media (Lonza). They were stimulated for 24 hours with IL-1β, IL-6, IFN-γ, osteopontin, or TNF-α, all at 10 ng/mL and purchased from PeproTech, or 10 ng/mL TGF-β1, 1.1 ng/mL TGF-β2, or 5.5 ng/mL TGF-β3, all three from R&D Systems. In inhibition experiments, cells were pre-incubated with the NF-kB inhibitor (10 uM) for 1 hour or transfected with a p65-targeted siRNA (50 nM; Santa Cruz Biotechnology, Santa Cruz, CA, cat. #sc-29410) or scrambled siRNA (Santa Cruz Biotechnology, cat. #sc-44230 [Scrambled B]). siRNA transfection was performed using JetPEI (Polyplus, Illkirch-Graffenstaden, France, cat. #101000053) according to the manufacturer’s protocol. The effect of laminal flow was tested using a Streamer Fluid Shear device (FlexCell International, Burlington, NC, USA). For this, 1 × 10^6^ cells were seeded in slides recommended by the manufacturer. After 24 hours, the confluent cell cultures were exposed to laminar flow for 6 hours (5±3 dynes/cm^2^) or static conditions, in the presence or absence of IL-1β (10 ng/mL). Cell passages 5 to 7 were used in all experiments.

### RNA Extraction and Real-Time PCR

RNA was isolated using the E.Z.N.A Total RNA kit (Omega Bio-tek, Norcross, GA; cat.# R6812-02) and cDNA synthesized with the High-Capacity cDNA Reverse Transcription kit (Applied Biosystems, Waltham, MA). Real-time PCR was performed on an ABI QuantStudio 3 Real-Time PCR System (Applied Biosystems) using the following TaqMan assays: *ACKR3*, Hs00664172_s1; *ADAM12*, Hs01106101_m1; *ASS1*, Hs00540723_m1; *COL1A1*, Hs00164004_m1; *COL4A1*, Hs00266237_m1; *COL8A1*, Hs00156669_m1; *CYP1B1*, Hs00164383_m1; *GAPDH*, Hs02786624_g1; *HAS1*, Hs00758053_m1; *HAS2*, Hs00193435_m1; *MMP2*, Hs01548727_m1; *NOS3*, Hs01574665_m1; *PDPN*, Hs00366766_m1; *PTGDS*, Hs00168748_m1; *PTGIS*, Hs00919949_m1; *PTGS2*, Hs00153133_m1; *PTHLH*, Hs00174969_m1; and *TAGLN*, Hs01038777_g1. Relative gene expression was determined using the ΔΔCT method and normalized with respect to *GAPDH*. Gene expression changes were reported as fold change compared to control conditions.

### Flow Cytometry

The identity of primary AVF fibroblasts was confirmed by flow cytometry for CD90 and PDGFRA. An aliquot of cells was incubated with 1 *μ*L of the LIVE/DEAD Fixable Dead Stain Kit (Invitrogen) at 4°C for 30 min. The rest of the sample was resuspended in FACS buffer (PBS supplemented with 2% FBS, 1 mM EDTA, 0.1% sodium azide) and incubated with 8 *μ*L of Human BD Fc Block (BD Biosciences, Franklin Lakes, NJ) at 4°C for 10 min. After washing and aliquoting into individual flow tubes, cells were labeled for 30 min with 2 *μ*L of the target antibodies (CD90 PE-Cy7, BD Biosciences cat. #561558; PDGFRA PE, BD Biosciences, cat. # 556002), including single-stain and no-stain controls. All tubes were washed with FACS buffer, followed by fixation with 2% PFA in PBS for 15 min at room temperature. Cells were washed again and stored at 4°C in FACS buffer overnight. Samples were read in a BD LSRFortessa Cell Analyzer and analyzed using FlowJo Software v10.10 (BD Biosciences).

### Histology and Immunofluorescence

For immunofluorescence, paraffin sections were rehydrated by serially immersing them in xylene, alcohol, and water, and antigens retrieved by boiling slides in 10 mM citrate buffer, pH 6.0 or Tris-EDTA buffer, pH 9.0 for 30 minutes (see table below). Sections were treated with 3% hydrogen peroxide and TNB Blocking Buffer (Akoya Biosciences, Marlborough, MA; cat. #FP1012), followed by primary antibodies diluted in TNB overnight at 4^ο^C. The table below indicates the dilution ratio of specific antibodies. Bound antibodies were detected with Alexa Fluor 546 goat anti-rabbit antibody (1:1000, #A11081; all secondary antibodies from Thermo Fisher Scientific), Alexa Fluor 633 goat anti-mouse antibody (1:1000, #A21052), or Alexa Fluor 633 goat anti-rabbit antibody (1:1000, #A21071) for 45 minutes. Sections were counter-stained with 300 nM DAPI solution (#D1306, Thermo Fisher Scientific) in PBS for 5 minutes, and mounted in fluorescence-compatible mounting medium (Abcam, Cambridge, United Kingdom; cat. #AB104135-1002). For signal amplification, after overnight incubation with primary antibodies, slides were incubated with biotinylated goat anti-mouse (1:1000, #OS02B; Oncogene Research, La Jolla, CA) antibody for 1 hour, then streptavidin HRP (1:1000, #P0397; DAKO, Santa Clara, CA) for 30 minutes, and finally amplified with a Tyramide Signal Amplification kit (1:50, #NEL700A001KT, Perkin Elmer, Waltham, MA). Amplification of the biotinylated antibody was detected through incubation with a streptavidin conjugated Alexa 546 secondary (1:1000, #S11225, Thermo Fisher Scientific) for 1 hour. Sections were examined in a Keyence All-in-One Fluorescence Microscope BZ-X800L and photographed using the Keyence BZ-X800 Viewer software (Keyence, Itasca, IL). Morphometry was measured in SMA stained sections using ImageJ (National Institutes of Health).

### In Situ Hybridization with Immunofluorescence

In situ hybridization (ISH) was performed using the RNAscope Detection Kit (Advanced Cell Diagnostics, Newark, CA) with the *POSTN*-specific probe (ACD-409181-C3), followed by immunofluorescence staining. Briefly, formalin-fixed paraffin-embedded tissue sections were deparaffinized in xylene, rehydrated through a graded ethanol series, and treated with hydrogen peroxide for 10 minutes to block endogenous peroxidase activity. Antigen retrieval was carried out using the RNAscope Target Retrieval Reagent at 95–100°C for 15 minutes, followed by incubation with RNAscope Protease Plus at 40°C for 30 minutes. The *POSTN* probe was hybridized to the tissue sections at 40°C for 2 hours in a humidity-controlled chamber. Signal amplification and detection were performed according to the manufacturer’s instructions, and sections were thoroughly washed in RNAscope Wash Buffer between steps. For immunofluorescence, tissues were blocked with TNB Blocking Buffer for 1 hour and incubated overnight at 4°C with anti-HAS1 antibody (Bioss, Woburn, MA; cat. #BS2946R). Sections were then washed and incubated with the fluorophore-conjugated secondary antibody Alexa 546 (Invitrogen, #A-11035) for 1 hour at room temperature. Slides were counterstained with DAPI to visualize nuclei, mounted with fluorescence-compatible mounting medium (Abcam, #AB104135-1002), and imaged using the Keyence All-in-One Fluorescence Microscope BZ-X800L.

### Statistical Analysis

Statistical analyses were performed using GraphPad Prism 10.1.1 (San Diego, CA). Normally distributed data were compared using t-tests and expressed as mean ± standard deviation (SD). If normality assumptions were not met, the Mann-Whitney test was used, and data were expressed as median and interquartile range (IQR). Comparisons of paired vein and AVF samples from the same patient were performed using paired t-tests or paired Wilcoxon signed-rank tests as appropriate.

| **TARGET** | **ANTIBODY** | **DILUTION** | **ANTIGEN RETRIEVAL** |
| --- | --- | --- | --- |
| KI67 | Abcam – ab833 | 1:50 | pH 6 |
| SMA | Cell Signaling – 1924SS | 1:100 | pH 6 |
| MYH11 | Sigma – M7786 | 1:100 | pH 6 |
| PDGFRA | Origene – TAB04956 | 1:100 + AMP* | pH 9 |
| CD68 | Abcam – ab213363 | 1:100 | pH 9 |
| CD31 | DAKO – M0823 | 1:50 | pH 9 |
| ACKR3 | Invitrogen – PA3-069 | 1:100 | pH 9 |

\*AMP: Amplification

